# AptaRender: an interactive drawing and styling tool for 2D nucleic acid structures

**DOI:** 10.64898/2026.11.12.766329

**Authors:** Yujie He, Yui Zhu, Hieu Tran, Hsin-Chih Yeh

## Abstract

Two-dimensional (2D) diagrams are widely used to communicate nucleic acid secondary structure, yet the path from a sequence and pairing model to an editable publication figure often requires either specialized software or substantial cleanup in a general-purpose graphics editor. AptaRender (v2.3.1, available at https://yjhe99.github.io/aptarender/) is a browser-based editor that converts a set of user-supplied DNA or RNA sequences and their dot-bracket structures into an interactive 2D drawing. The software separates structure specification from visual composition: it does not predict folding, but provides direct control over the layout, nucleotide styling, base pair styling, numbering and export. AptaRender supports simultaneously working with multiple sequences, customized per-nucleotide sizing and coloring with customized styles, manual branching and motif repositioning, constrained rotation of helices around loop rings, and visualization of a user defined hybridization region between two strands. Drawings are exported as editable SVG or high-resolution PNG. The application is implemented as a single dependency-free HTML file that runs entirely in the browser and stores working state locally. This design makes the tool easy to distribute and suitable for rapid preparation of figures for manuscripts, presentations, and exploratory communication. AptaRender complements prediction and template-based visualization software by concentrating on lightweight, local, art directed figure construction. Across eight test cases, AptaRender reliably generated accurate, information-rich 2D nucleic acid structures and exported them in PNG and SVG formats, although specialized annotations and additional symbols required external editing.

## INTRODUCTION

Secondary structure diagrams provide a compact representation of base pairing and help researchers relate sequence features to nucleic acid function. However, the challenges with visualization are distinct from structure prediction. Prediction methods infer a pairing model, whereas visualization software must translate that model into a legible arrangement of stems, loops, junctions, termini, labels, and annotations. The resulting figure often needs further editing before it can communicate on a particular experiment or mechanism.

Several established programs address parts of this workflow. VARNA combines automated layouts with interactive editing and annotation^1^. Forna uses a force-directed layout in a web interface and supports interactive exploration^2^. R2R focuses on aesthetic consensus diagrams^3^, RNApuzzler emphasizes intersection free outerplanar layouts^4^, and R2DT produces recognizable template-based layouts at scale^5,6^. RNA2Drawer and RNAcanvas provide geometrically controlled and highly editable drawings^7,8^. However, these tools do not provide the lightweight workflow for rapidly styling and refining user-supplied structures.

To address this need, we developed AptaRender as a lightweight browser-based editor, for turning a supplied DNA or RNA pairing model into a clear and editable figure. The application favors immediate browser access (available at https://yjhe99.github.io/aptarender/), visible controls, local data handling, and editable vector export. The current release substantially extends the initial public version with multiple sequence management, structured selection and movement, data driven coloring, two strand hybridization diagrams, constrained loop centered helix rotation, and a unified rendering and export path. This article describes the design, implementation, capabilities, and appropriate scope of AptaRender. AptaRender overlaps with several established programs but differs in scope and packaging. It is best understood as a lightweight figure editor for supplied, nested secondary structures. It does not replace tools designed for folding, pseudoknot layout, consensus diagrams, standardized templates, tertiary interaction notation, or large RNA families.

**Table 1.** Comparison of AptaRender relative to representative published tools.

| Tool | Primary emphasis | Interactive web editing | Selected features |
|---|---|---|---|
| AptaRender | Art directed figures from supplied nested structures | Yes | - a single lightweight browser app<br>- Customizable per-base styling<br>- Enables two-strand complex schematic |
| VARNA<sup>1</sup> | Automated drawing, editing, and annotation | Yes | - Enables variable exported file formats<br>- Utilizes multiple layout algorithms |
| Forna<sup>2</sup> | Interactive force directed diagrams | Yes | Enables pseudoknot visualization |
| R2R<sup>3</sup> | Aesthetic consensus diagrams | No | Alignment-driven consensus annotation |
| RNApuzzler<sup>4</sup> | Automated outerplanar layout | No | Intersection-free drawings for supported structures |
| R2DT<sup>5,6</sup> | Recognizable template-based layouts | Via linked editors | Contains multiple layout templates |
| RNAcanvas<sup>8</sup> | Detailed interactive nucleic acid drawing | Yes | Allows for extensive graphical editing and interaction notation |
| RNA2Drawer<sup>7</sup> | Geometrically constrained drawing and graphical editing | Yes | Highlights complementary subsequences |

## DESIGN GOALS

AptaRender was designed around five requirements. First, a user should be able to open the application and draw a structure without installing software or learning a scripting language. Second, the application should treat the supplied pairing model as authoritative and avoid implying that a visually pleasing layout is a structural prediction. Third, users should be able to adjust the figure at the level of individual nucleotides, structural parts, or complete branches.

Fourth, the on-screen drawing and exported figure should follow the same rendering logic. Fifth, sequence and styling data should remain in the browser unless the user explicitly exports a file.

## SOFTWARE IMPLEMENTATION

### Input model and parsing

The application accepts DNA or RNA sequences containing A, U, T, G, C, or N and a structure string of equal length. Periods and selected punctuation denote unpaired positions. Parentheses denote canonical pairs, while square, curly, or angle brackets mark pairs that are drawn with dashed rungs to distinguish non-Watson-Crick interactions. Brackets must be properly nested; crossing pseudoknots are not supported. The parser returns a pair table, flags noncanonical pairs, and reports length mismatches, invalid characters, unbalanced delimiters, and mismatched bracket types with position specific messages.

### Layout and scene generation

A radial layout engine places paired stems and enclosed loops while treating exterior unpaired segments as free tails or linkers. Hairpin, internal, and multibranch loops are arranged around circular arcs. Exterior segments can be rendered with natural, flattened, L shaped, or perpendicular tail styles. Manual offsets are stored separately from automatically generated coordinates so users can reset edits without changing the sequence or structure.

A single scene generator creates the ordered graphical elements used by both the interactive canvas and export functions. The scene contains duplex rungs, intramolecular pair rungs, backbones, nucleotide circles, position labels, and nucleotide letters. Interactive hit targets and selection indicators are added only for the canvas. The same noninteractive scene is serialized for SVG export or rasterized for PNG export, reducing discrepancies between the editor and downloaded figures.

### Interaction and state

Users can select individual nucleotides, add to a selection, box select, or select a structural part inferred from the pair table. Structural parts include a stacked helix, the unpaired residues of a loop ring, and an exterior unpaired run. Dragging can move a branch, a selected group, one inferred part, or a single nucleotide. Selected bases can also be rotated about their centroid or attachment point. A loop locking operation allows an attached helix to swing around a loop while preserving the ring radius and redistributing unpaired residues between neighboring helices.

The application maintains up to 100 undo states. Multiple sequences and two strand complexes can be stored in the same browser session. Working data are serialized to browser local storage, so no server upload is required for normal use. The full application is distributed as one HTML file containing its markup, style rules, and JavaScript, with no runtime framework or external library.

## VISUALIZATION AND ANNOTATION CAPABILITIES

AptaRender provides controls for nucleotide letters, nucleotide circles, backbone lines, pair rungs, position numbers, background, and export. Letter colors may be assigned globally, by nucleotide identity, by selection, or from a user supplied text file. The same file mechanism can set circle fills and accepts positions, ranges, optional strand prefixes, optional base identity checks, and CSS color values. Per nucleotide scaling emphasizes selected sites without changing the global font size.

Position labels are placed in available space around each residue. Candidate positions are tested against nucleotide circles, line segments, and previously placed labels. When a label cannot remain close to its nucleotide, the drawing adds a leader line. Noncanonical pairs are represented by dashed rungs. Individual intramolecular rungs can be recolored in single sequence view, and the interstrand rungs of a complex have a separate color.

## TWO STRAND COMPLEXES

A partially hybridized nucleic acid complex is a common representation in multiple fields including functional nucleic acids^9–12^, DNA nanotechnology^13–16^, synthetic biology^17–19^, structural biology^20,21^, gene editing^22–24^ and RNA biology^25–27^. Such complexes join contiguous regions from two stored sequences in an antiparallel orientation, yet existing tools do not support their direct rendering. AptaRender addresses this gap by enabling users to render and easily refine two-strand complexes (**Fig. 2**). In Replace mode (**Fig. 2B**), intramolecular pairs that touch or span the designated hybridization region are removed and the selected regions are drawn as a straight duplex with the remaining structure attached at its ends. Replace mode also allows multiple hybrid regions which is typical in a bubble construct^28^ or a trans-cleavage DNAzyme^29^. In Side-by-side mode (**Fig. 2C**), the layout and intramolecular pairs of strand A are retained, while the selected region of strand B is positioned beside it. These modes support schematic communication of a proposed or experimentally defined interaction. They do not predict hybridization or test whether the resulting pairing is physically realizable.

**Figure 1.**
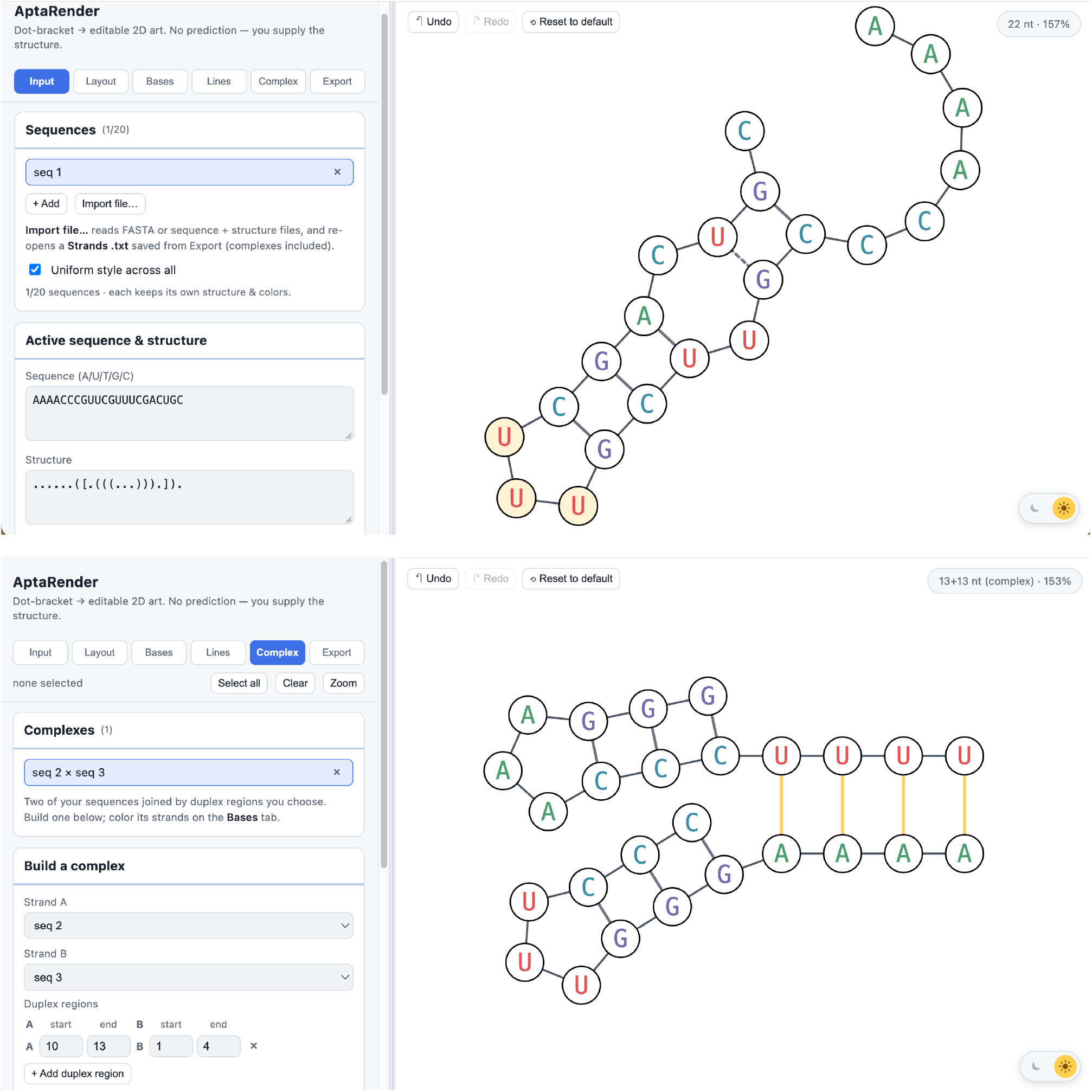
Representative AptaRender views from the user tutorial. The interface combines tabbed controls with an SVG canvas and supports single sequence styling, manual layout adjustment, and two strand complex figures.

**Figure 2.**
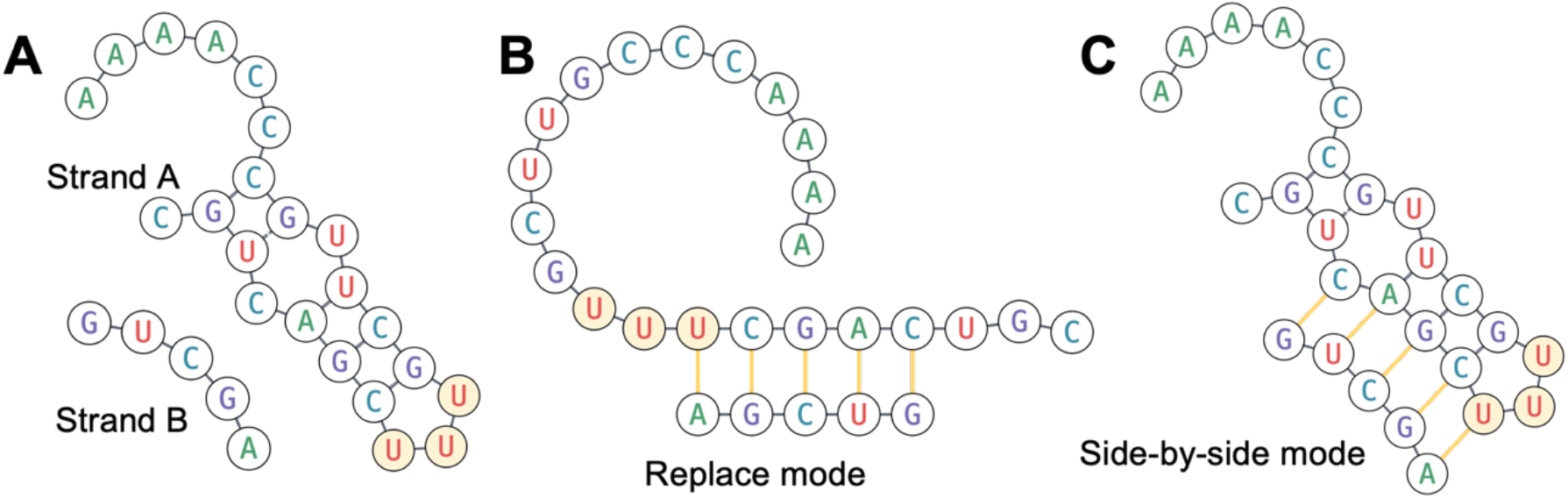
Representative visualizations of a two-strand complex generated using AptaRender. (**A**) Individual strands A and B before complex formation. (**B**) Strands A and B visualized as a complex in Replace mode. (**C**) Strands A and B visualized as a complex in Side-by-side mode. The sequences shown here are for illustrative purposes only and have no biological or chemical significance.

## REPRESENTATIVE WORKFLOWS

Users can access the tool at https://yjhe99.github.io/aptarender/ before starting a workflow. Four workflows illustrate the intended use of the software. A user can prepare a publication figure by supplying a sequence and dot bracket structure, applying a nucleotide palette, enabling circles and numbering, adjusting crowded branches, and exporting SVG. A binding site or chemically probed region can be emphasized by selecting residues or importing a position color file. A multibranch structure can be clarified by rotating a selected branch about its attachment point or swinging a helix around a locked loop ring. After that, two stored strands can be combined to depict a specified interaction region and styled with a distinct duplex rung color. Currently, AptaRender (v2.3.1) does not support customized annotations or shapes such as arrows and additional labels. These can be added outside the software after exporting.

## TECHNICAL EVALUATION

### Test set

The test set comprised eight literature-derived cases with individual strand lengths ranging from 11 to 346 nucleotides (**Fig. 3**). The examples were selected to represent structures including hairpins (test cases 1, 7-2 and 8-1), internal loops (test cases 3), multibranch junctions (test cases 2, 4 and 5), free strand or long free tails (test cases 7-3, 8-2 and 8-3), noncanonical base pairs (test cases 4 and 7-2), and two-strand complexes (test cases 3, 6 and 7-1) with various visuals applied including color-coding and strand curving. The selected cases also covered a wide range of nucleic acid scaffolds, including aptamers (test cases 2, 4, 7 and 8), riboswitches (test case 4), DNAzymes (test case 5), NCBs (nanocluster beacons, test cases 1 and 6) and unit of nanostructures (test case 3). Here since test cases 7 and 8 has multiple drawings, individual one was referred to as -1, -2, and -3, respectively. For example, test case 7-1 refers to the first structure in test case 7, i.e., the partial DNA duplex (**Fig. 3N, left**). Sequences and base-pairing models were obtained from the corresponding published figures and cited sources^**9**,**15**,**30–35**^. Each structure was independently redrawn with AptaRender (v2.3.1) to benchmark the reproducibility. The eight structures, their lengths, structural classes, and test environments are summarized in **Table 2**.

**Table 2.** Technical evaluation of representative nucleic acid structures. nt, nucleotide; R-FAP, RNA-based fluorogenic aptamer; MgZ, magnesium-activated DNAzyme; 3WJ, three-way junction; NCB, nanocluster beacon; AgNC, silver nanocluster; F, fluorophore (label); Q, quencher (label).

| **#** | **Name [ref]** | **Length (nt)** | **Structure class** | **Output** | **Platform** | **Manual additions** | **Notes** |
|---|---|---|---|---|---|---|---|
| **1** | rSubak [30] | 37 | hairpin | PNG/SVG: successful | Chrome on macOS | 1. Text annotations<br>2. Star labels | Shading replaced by colored circles |
| **2** | Lettuce [31] | 67 | Multiloop aptamer | PNG/SVG: successful | Chrome on macOS | 1. Text annotations<br>2. Custom nucleotide numbering | Shadings replaced by colored circles |
| **3** | Internal-loop RNA [32] | 13+8 | Partial RNA duplex | PNG/SVG: successful | Chrome on macOS | Dot labels |  |
| **4** | R-FAP-based glycine sensor [34] | 346 | Highly branched riboswitch-aptamer | PNG/SVG: successful, with/without nt labels | Chrome on macOS | 1. Text annotations<br>2. F symbols<br>3. Bond-line format | AptaRender shows more structural detail than the simplified reference |
| **5** | MgZ in cis [33] | 83 | 3WJ DNAzyme | PNG/SVG: successful | Edge on Windows | 1. Text annotations<br>2. Arrow labels<br>3. A-T/G-C pair styling<br>4. Ribonucleotide labels | Modified dT shown as T rather than F/Q |
| **6** | NCB [15] | 48+48 | Partial DNA duplex | PNG/SVG: successful, with/without nt labels | Opera on macOS | 1. Text annotations<br>2. AgNC symbols<br>3. Arrow labels |  |
| **7** | Strand-displacement aptasensor [9] | 41+11 | Partial DNA duplex | PNG/SVG: successful | Chrome on macOS | 1. Text annotations<br>2. F/Q/ protein symbols<br>3. Arrow label |  |
|  |  | 41 | Aptamer hairpin |  |  |  |  |
|  |  | 11 | Free strand |  |  |  |  |
| **8** | Aptamer Beacon [35] | 53 | Hairpin | PNG/SVG: successful, with/without nt labels | Chrome on macOS | 1. Text annotations<br>2. F/Q/ligand symbols |  |
|  |  |  | Free strand | PNG/SVG: successful |  |  |  |
|  |  |  | Multiloop with a free tail | PNG/SVG: successful, with/without nt labels |  |  |  |

**Figure 3.**
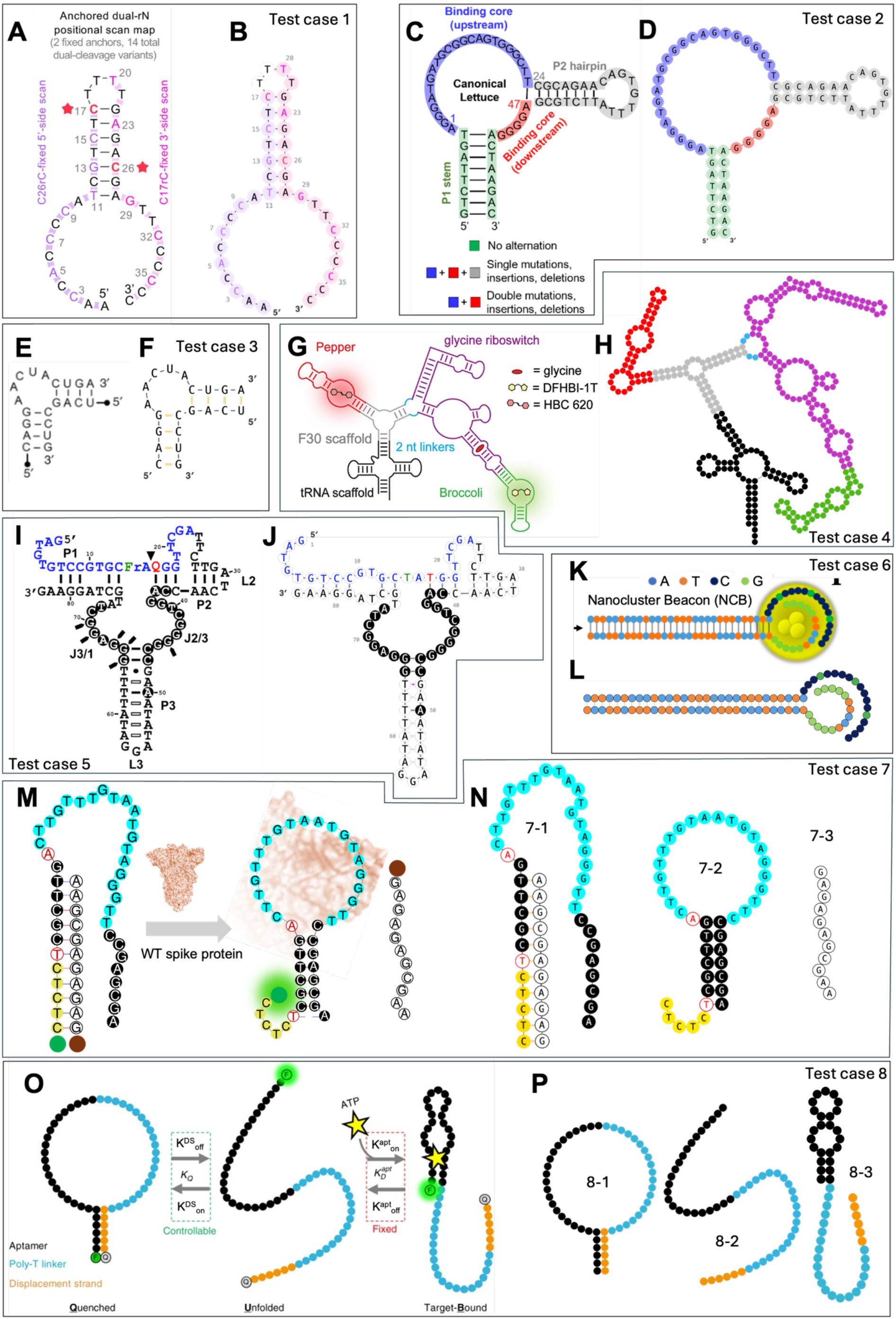
Test cases reproduced with AptaRender 2.2.4. Reference figures and reproduced figures are arranged side by side. Reference structures used to evaluate AptaRender: (**A**) Test case 1, rSubak hairpin structure. Reprinted in part from Fig. 2a of Kim *et al*.^30^. Copyright (2026) the authors/funder, licensed under a Creative Commons Attribution-NonCommercial 4.0 International License. (**B**) rSubak independently reproduced with AptaRender. (**C**) Test case 2, Lettuce fluorogenic aptamer structure. Reprinted in part from Fig. 1h of Kuo *et al*.^31^ under the author-reuse terms of ACS Publications. Copyright (2026) American Chemical Society. (**D**) Lettuce independently reproduced with AptaRender. (**E**) Test case 3, HCV internal-loop RNA structure. Reprinted in part from Fig. 1a of Monferrer *et al*.^32^. Copyright (2019) The Authors, licensed under a Creative Commons Attribution 4.0 International License. (**F**) The HCV internal-loop structure independently reproduced with AptaRender. (**G**) Test case 4, RNA fluorogenic aptamer-based glycine sensor structure. Reprinted in part from Fig. 4a of Bodin and Hammond^34^. Copyright (2025) The Authors, licensed under a Creative Commons Attribution-NonCommercial 4.0 International License. (**H**) Glycine sensor independently reproduced with AptaRender. (**I**) Test case 5, MgZ three-way-junction DNAzyme structure. Reprinted in part from Fig. 1b of Chiuman and Li^33^. Copyright (2007) Chiuman and Li, licensed under a Creative Commons Attribution License. (**J**) MgZ independently reproduced with AptaRender. (**K**) Test case 6, NanoCluster beacon (NCB) two-strand complex. Reprinted in part from Fig. 1a of Kuo *et al*.^15^ under Wiley’s author-reuse policy. Copyright (2022) Wiley-VCH GmbH. (**L**) NCB independently reproduced with AptaRender. (**M**) Test case 7, structure-switching SARS-CoV-2 DNA aptasensor. Reprinted in part from Fig. 7a of He *et al*.^9^ under the author-reuse terms of ACS Publications. Copyright (2026) American Chemical Society. (**N**) The aptasensor independently reproduced with AptaRender. (**O**) Test case 8, intramolecular aptamer switch. Reprinted from Fig. 1 of Wilson *et al*.^35^. Copyright (2019) The Authors, licensed under a Creative Commons Attribution 4.0 International License. (**P**) The intramolecular aptamer switch independently reproduced with AptaRender. Panels **C, K**, and **M** are excluded from any Creative Commons license applied to this manuscript; reuse of those panels remains subject to the respective copyright holders’ terms.

### Evaluation criteria

Each test case was evaluated for successful parsing, rendering, and export in SVG and PNG formats. Visual quality was assessed by comparing the original structure with the rendered one and identifying unsupported features which require post-processing with external tools (e.g., adding quenchers and labels to the construct). Error messages were documented. **Table 2** summarizes the structure lengths and classes, test environments, export outcomes, and features requiring manual addition.

### Results

All eight test cases listed in **Table 2** were successfully processed and the resulting visualizations were exported in both PNG and SVG formats without errors. Although AptaRender generated the core structural representations, every case required some degree of post-processing to fully reproduce the corresponding reference figure. These edits included the addition or refinement of text elements, such as section names, terminal labels, and legends, as well as graphical annotations, including arrows, chemical modifications, and their associated labels. These elements are not currently supported by AptaRender and therefore had to be added using external graphics-editing software. In addition, the current version of AptaRender (v2.3.1) does not support the simplified bond-line representation used in test case 4, as this type of schematic is better suited to general-purpose illustration software than to structure-rendering software.

Test cases 1, 2, 3, 6, 7, and 8 were reproduced accurately because their reference schematics depicted structurally correct 2D conformations, which were consistent with predictions generated by established nucleic acid structure-prediction software (**Fig. 3A-F, K-N**). Test case 4 produced a secondary structure that differed slightly from the reference schematic (**Fig. 3G, H**). This arose from the fact that the source schematic was a simplified representation that did not fully capture the predicted scaffold architecture. Consequently, the AptaRender visualization provided a more detailed and structurally accurate representation of the underlying nucleic acid scaffold (**Fig. 3H**). For test case 5, the apparent pairing between nucleotide 37A and the quencher reflects base pairing between 37A and the underlying thymidine (dT) to which the quencher is attached. In the AptaRender reconstruction, the modified nucleotide was represented as T rather than by the fluorophore/quencher labels (F and Q) with the original color code, thereby preserving the original sequence and correctly depicting the base-pairing interaction (**Fig. 3I, J**). For reference schematics that omitted base identities, nucleotide-labeled versions were also generated to demonstrate AptaRender’s distinctive ability to produce clear, information-rich structural visualizations with minimal additional effort. These labeled renderings remained visually clear and well organized (**Supplementary Figs. 1-2**). **Table 2** provides a case-by-case summary of these outcomes and the additional editing required. Overall, these results demonstrate that AptaRender can reliably generate and export the principal 2D structures, while specialized labels and nonstructural annotations may require additional manual editing.

## DISCUSSION

AptaRender addresses the part of the visualization workflow that often remains after a pairing model has already been chosen: making a diagram readable, emphasizing experimental observations, and exporting artwork that can be edited or placed directly in a figure. This browser-only implementation increases the incentives to trying the software and makes the application easy to archive and deploy. Local storage is useful for unpublished sequences because routine editing does not require data transmission, although users must export important work before clearing browser data or moving to another computer.

The design also imposes clear limits. The parser accepts only nested pairings and therefore cannot represent crossing pseudoknots. Complex view supports two strands and one contiguous interstrand duplex. Side-by-side mode can depict a strand A nucleotide with both an intramolecular and interstrand bond, which is useful schematically but may be physically impossible. The software does not calculate thermodynamic stability, predict structures, validate pair identities, or infer hybridization. These constraints should remain visible in the interface and documentation so visual output is not mistaken for computational or experimental evidence.

Future development should be guided by reproducible user needs. Priority areas include a portable project format, broader interaction notation, accessibility testing, automated regression examples, simplified GUI, and optional interoperability with a community exchange format such as the RNA 2D JSON Schema described by R2DT 2.0^5^. Support for pseudoknots or additional strands would require changes to both the data model and layout engine and should not be treated as a cosmetic extension.

## Supporting information

Exported files from technical evaluations

User tutorial

## ACKNOWLEDGEMENTS

We thank Sohyun Kim, Anh-Thu Nguyen, Stefen Stangherlin, Jesse Donovan and Gabriel Alejandro Romero Agosto for their valuable feedback and suggestions on the first version of AptaRender.

## Author Contributions

Y. H. and H.-C. Y. discussed and defined the project. Y. H. developed the first version of AptaRender. H. T. and Y. Z. updated AptaRender. Y. H., Y. Z., and H. T. performed the technical evaluation. Y. H. wrote the manuscript with the help of all the co-authors. H. T. and Y. Z. reviewed the manuscript. H.-C. Y. supervised the project.

## Data and materials availability

AptaRender is available at https://yjhe99.github.io/aptarender/. The version of AptaRender evaluated in this study (v2.3.1) is archived in Zenodo at https://doi.org/10.5281/zenodo.24185163. The current source code and subsequent releases are available at https://github.com/yjhe99/aptarender.

## Supplementary Materials

Sequences and corresponding dot-bracket annotations used in this report, selected test cases with nucleotide labels; user tutorial; all exported .svg and .png files from the technical evaluation.

## References

(1) Darty, K.; Denise, A.; Ponty, Y. VARNA: Interactive Drawing and Editing of the RNA Secondary Structure. Bioinformatics 2009, 25 (15), 1974–1975. 10.1093/bioinformatics/btp250

(2) Kerpedjiev, P.; Hammer, S.; Hofacker, I. L. Forna (Force-Directed RNA): Simple and Effective Online RNA Secondary Structure Diagrams. Bioinformatics 2015, 31 (20), 3377–3379. 10.1093/bioinformatics/btv372

(3) Weinberg, Z.; Breaker, R. R. R2R - Software to Speed the Depiction of Aesthetic Consensus RNA Secondary Structures. BMC Bioinformatics 2011, 12, 3. 10.1186/1471-2105-12-3

(4) Wiegreffe, D.; Alexander, D.; Stadler, P. F.; Zeckzer, D. RNApuzzler: Efficient Outerplanar Drawing of RNA-Secondary Structures. Bioinformatics 2019, 35 (8), 1342–1349. 10.1093/bioinformatics/bty817

(5) McCann, H.; Meade, C. D.; Williams, L. D.; Petrov, A. S.; Johnson, P. Z.; Simon, A. E.; Hoksza, D.; Nawrocki, E. P.; Chan, P. P.; Lowe, T. M.; Ribas, C. E.; Sweeney, B. A.; Madeira, F.; Anyango, S.; Appasamy, S. D.; Deshpande, M.; Varadi, M.; Velankar, S.; Zirbel, C. L.; Naiden, A.; Jossinet, F.; Petrov, A. I. R2DT: A Comprehensive Platform for Visualizing RNA Secondary Structure. Nucleic Acids Res 2025, 53 (4), gkaf032. 10.1093/nar/gkaf032

(6) Sweeney, B. A.; Hoksza, D.; Nawrocki, E. P.; Ribas, C. E.; Madeira, F.; Cannone, J. J.; Gutell, R.; Maddala, A.; Meade, C. D.; Williams, L. D.; Petrov, A. S.; Chan, P. P.; Lowe, T. M.; Finn, R. D.; Petrov, A. I. R2DT Is a Framework for Predicting and Visualising RNA Secondary Structure Using Templates. Nat Commun 2021, 12 (1), 3494. 10.1038/s41467-021-23555-5

(7) Johnson, P. Z.; Kasprzak, W. K.; Shapiro, B. A.; Simon, A. E. RNA2Drawer: Geometrically Strict Drawing of Nucleic Acid Structures with Graphical Structure Editing and Highlighting of Complementary Subsequences. RNA Biol 2019, 16 (12), 1667–1671. 10.1080/15476286.2019.1659081

(8) Johnson, P. Z.; Simon, A. E. RNAcanvas: Interactive Drawing and Exploration of Nucleic Acid Structures. Nucleic Acids Res 2023, 51 (W1), W501–W508. 10.1093/nar/gkad302

(9) He, Y.; Yang, Z.; Kuo, Y.-A.; Wu, Y.; Fonseca-Albert, D.; Le, K. K.; Guo, J.; Wang, Y.; Nguyen, A.-T.; Chen, Y.-I.; Kim, S.; Chen, W.-R.; Seifi, S.; Hong, S.; Nguyen, T. D.; Chen, Y.; Ren, P.; Lu, Y.; Yeh, H.-C. High-Throughput On-Chip Screening Enables Rapid Adaptation of DNA Aptamers to SARS-CoV-2 Evolution. ACS Nano 2026, 20 (34), 23941–23957. 10.1021/acsnano.6c08330

(10) Hariri, A. A.; Cartwright, A. P.; Dory, C.; Gidi, Y.; Yee, S.; Thompson, I. A. P.; Fu, K. X.; Yang, K.; Wu, D.; Maganzini, N.; Feagin, T.; Young, B. E.; Afshar, B. H.; Eisenstein, M.; Digonnet, M. J. F.; Vuckovic, J.; Soh, H. T. Modular Aptamer Switches for the Continuous Optical Detection of Small-Molecule Analytes in Complex Media. Advanced Materials 2024, 36 (1), 2304410. 10.1002/adma.202304410

(11) Stangherlin, S.; Ding, Y.; Liu, J. Dissociation Constant (Kd) Measurement for Small-Molecule Binding Aptamers: Homogeneous Assay Methods and Critical Evaluations. Small Methods 2025, 9 (6), 2401572. 10.1002/smtd.202401572

(12) Yang, Z.; Shao, X.; Wu, Y.; Roy, A.; Garcia, E.; Farrell, A.; Pradhan, S.; Guo, W.; Gan, H.; Korkmaz, Z.; Adams, E.; Lu, Y. Decoding Potassium Homeostasis in Cancer Metastasis and Drug Resistance: Insights from a Highly Selective DNAzyme-Based Intracellular K+ Sensor. J Am Chem Soc 2025, 147 (21), 18074–18087. 10.1021/jacs.5c03781

(13) Zhang, Y.; McMullen, A.; Pontani, L.-L.; He, X.; Sha, R.; Seeman, N. C.; Brujic, J.; Chaikin, P. M. Sequential Self-Assembly of DNA Functionalized Droplets. Nat Commun 2017, 8 (1), 21. 10.1038/s41467-017-00070-0

(14) Green, L. N.; Subramanian, H. K. K.; Mardanlou, V.; Kim, J.; Hariadi, R. F.; Franco, E. Autonomous Dynamic Control of DNA Nanostructure Self-Assembly. Nat. Chem. 2019, 11 (6), 510–520. 10.1038/s41557-019-0251-8

(15) Kuo, Y.-A.; Jung, C.; Chen, Y.-A.; Kuo, H.-C.; Zhao, O. S.; Nguyen, T. D.; Rybarski, J. R.; Hong, S.; Chen, Y.-I.; Wylie, D. C.; Hawkins, J. A.; Walker, J. N.; Shields, S. W. J.; Brodbelt, J. S.; Petty, J. T.; Finkelstein, I. J.; Yeh, H.-C. Massively Parallel Selection of NanoCluster Beacons. Advanced Materials 2022, 34 (41), 2204957. 10.1002/adma.202204957

(16) Kang, H.; Yang, Y.; Wei, B. Synthetic Molecular Switches Driven by DNA-Modifying Enzymes. Nat Commun 2024, 15 (1), 3781. 10.1038/s41467-024-47742-2

(17) Takahashi, M. K.; Tan, X.; Dy, A. J.; Braff, D.; Akana, R. T.; Furuta, Y.; Donghia, N.; Ananthakrishnan, A.; Collins, J. J. A Low-Cost Paper-Based Synthetic Biology Platform for Analyzing Gut Microbiota and Host Biomarkers. Nat Commun 2018, 9 (1), 3347. 10.1038/s41467-018-05864-4

(18) Kim, J.; Zhou, Y.; Carlson, P. D.; Teichmann, M.; Chaudhary, S.; Simmel, F. C.; Silver, P. A.; Collins, J. J.; Lucks, J. B.; Yin, P.; Green, A. A. De Novo-Designed Translation-Repressing Riboregulators for Multi-Input Cellular Logic. Nat Chem Biol 2019, 15 (12), 1173–1182. 10.1038/s41589-019-0388-1

(19) Zhao, E. M.; Mao, A. S.; de Puig, H.; Zhang, K.; Tippens, N. D.; Tan, X.; Ran, F. A.; Han, I.; Nguyen, P. Q.; Chory, E. J.; Hua, T. Y.; Ramesh, P.; Thompson, D. B.; Oh, C. Y.; Zigon, E. S.; English, M. A.; Collins, J. J. RNA-Responsive Elements for Eukaryotic Translational Control. Nat Biotechnol 2022, 40 (4), 539–545. 10.1038/s41587-021-01068-2

(20) Fang, C.; Huang, K.; Wu, X.; Zhang, H.; Gu, Z.; Wang, J.; Zhang, Y. Transcription Elongation of the Plant RNA Polymerase IV Is Prone to Backtracking. Sci Adv 10 (34), eadq3087. 10.1126/sciadv.adq3087

(21) Zhao, D.; Liu, W.; Chen, K.; Wu, Z.; Yang, H.; Xu, Y. Structure of the Human RNA Polymerase I Elongation Complex. Cell Discov 2021, 7 (1), 97. 10.1038/s41421-021-00335-5

(22) Pacesa, M.; Loeff, L.; Querques, I.; Muckenfuss, L. M.; Sawicka, M.; Jinek, M. R-Loop Formation and Conformational Activation Mechanisms of Cas9. Nature 2022, 609 (7925), 191–196. 10.1038/s41586-022-05114-0

(23) Liu, M.-S.; Gong, S.; Yu, H.-H.; Jung, K.; Johnson, K. A.; Taylor, D. W. Engineered CRISPR/Cas9 Enzymes Improve Discrimination by Slowing DNA Cleavage to Allow Release of off-Target DNA. Nat Commun 2020, 11 (1), 3576. 10.1038/s41467-020-17411-1

(24) Coelho, M. A.; De Braekeleer, E.; Firth, M.; Bista, M.; Lukasiak, S.; Cuomo, M. E.; Taylor, B. J. M. CRISPR GUARD Protects Off-Target Sites from Cas9 Nuclease Activity Using Short Guide RNAs. Nat Commun 2020, 11 (1), 4132. 10.1038/s41467-020-17952-5

(25) Vainberg Slutskin, I.; Weingarten-Gabbay, S.; Nir, R.; Weinberger, A.; Segal, E. Unraveling the Determinants of microRNA Mediated Regulation Using a Massively Parallel Reporter Assay. Nat Commun 2018, 9 (1), 529. 10.1038/s41467-018-02980-z

(26) Meng, L.; Liu, C.; Lü, J.; Zhao, Q.; Deng, S.; Wang, G.; Qiao, J.; Zhang, C.; Zhen, L.; Lu, Y.; Li, W.; Zhang, Y.; Pestell, R. G.; Fan, H.; Chen, Y.-H.; Liu, Z.; Yu, Z. Small RNA Zippers Lock miRNA Molecules and Block miRNA Function in Mammalian Cells. Nat Commun 2017, 8 (1), 13964. 10.1038/ncomms13964

(27) Hiers, N. M.; Li, L.; Li, T.; Sheng, P.; Wang, Y.; Traugot, C. M.; Yao, M.; Xie, M. An Endogenous Cluster of Target-Directed microRNA Degradation Sites Induces Decay of Distinct microRNA Families. Cell Rep 2025, 44 (9), 116162. 10.1016/j.celrep.2025.116162

(28) You, L.; Omollo, E. O.; Yu, C.; Mooney, R. A.; Shi, J.; Shen, L.; Wu, X.; Wen, A.; He, D.; Zeng, Y.; Feng, Y.; Landick, R.; Zhang, Y. Structural Basis for Intrinsic Transcription Termination. Nature 2023, 613 (7945), 783–789. 10.1038/s41586-022-05604-1

(29) Wu, Y.; Torabi, S.-F.; J. Lake, R.; Hong, S.; Yu, Z.; Wu, P.; Yang, Z.; Nelson, K.; Guo, W.; T. Pawel, G.; Stappen, J. V.; Shao, X.; M. Mirica, L.; Lu, Y. Simultaneous Fe2+/Fe3+ Imaging Shows Fe3+ over Fe2+ Enrichment in Alzheimer’s Disease Mouse Brain. Science Advances 2023. 10.1126/sciadv.ade7622

(30) Kim, S.; Hong, S.; Walker, J. N.; He, Y.; Nguyen, A.-T.; Chen, W.-R.; Seifi, S.; Chen, Y.-I.; Kuo, Y.-A.; Brodbelt, J. S.; Yeh, H.-C. Mechanistic Insights into the Color Transformation of a Non-FRET Substrate for RNase Activity Detection. bioRxiv March 13, 2026, p 2026.03.11.710959. 10.64898/2026.03.11.710959

(31) Kuo, Y.-A.; Chen, Y.-I.; Siraj, N.; He, Y.; Yang, Z.; Wang, Y.; Batchelder-Schwab, E. J.; Korkmaz, Z.; Yonas, S.; Nguyen, T. D.; Hong, S.; Nguyen, A.-T.; Kim, S.; Seifi, S.; Fan, P.-H.; Wu, Y.; Liu, H.-W.; Lu, Y.; Ren, P.; Mao, C.; Yeh, H.-C. Fluorogenic Aptamer Optimization on a Massively Parallel Sequencing Platform. ACS Sens. 2026, 11 (5), 3587–3600. 10.1021/acssensors.5c04046

(32) Monferrer, A.; Zhang, D.; Lushnikov, A. J.; Hermann, T. Versatile Kit of Robust Nanoshapes Self-Assembling from RNA and DNA Modules. Nat Commun 2019, 10 (1), 608. 10.1038/s41467-019-08521-6

(33) Chiuman, W.; Li, Y. Simple Fluorescent Sensors Engineered with Catalytic DNA ‘MgZ’ Based on a Non-Classic Allosteric Design. PLOS ONE 2007, 2 (11), e1224. 10.1371/journal.pone.0001224

(34) Bodin, M. R.; Hammond, M. C. Visualizing Intracellular Glycine with Two-Dye and Single-Dye Ratiometric RNA-Based Sensors. Nucleic Acids Res 2025, 53 (17), gkaf839. 10.1093/nar/gkaf839

(35) Wilson, B. D.; Hariri, A. A.; Thompson, I. A. P.; Eisenstein, M.; Soh, H. T. Independent Control of the Thermodynamic and Kinetic Properties of Aptamer Switches. Nat Commun 2019, 10 (1), 5079. 10.1038/s41467-019-13137-x

