## Supplementary Materials for "AptaRender: an interactive drawing and styling tool for 2D nucleic acid structures"

**Table S1. Sequences and their corresponding dot-bracket annotations used in this study**

| Name | Sequence / structure |
|---|---|
| Visualization only | |
| Main text **Fig. 1** top; **Fig. 2**-Strand A | AAAACCCGUUCGUUUCGACUGC |
| | ......([.(((...))).]). |
| Main text **Fig. 1** bottom-1 | GGGAAACCCUUUU |
| | (((...))).... |
| Main text **Fig. 1** bottom-2 | AAAAGGGUUUCCC |
| | ....(((...))) |
| Main text **Fig. 2**-Strand B | GUCGA |
| | ..... |
| Test cases | |
| Test case 1 (rSubak)[1] | AACCACCCCATCGTCTCTTTTGAGACGAGTTCCCCCC |
| | ..........(((((((....))))))))......... |
| Test case 2 (Lettuce)[2] | GTCTTAGTAGGGATGATGCGGCAGTGGGCTTCGCAG AACAGTGTTTATTCTGCGAGGGGACTAAGAC |
| | ((((((((.......................(((((((.........))))))).....)))))))) |
| Test case 3 (internal loop)-1[3] | CAGGAACUACUGA |
| | ............. |
| Test case 3 (internal loop)-2[3] | UCAGCCUG |
| | ........ |
| Test case 4 (R-FAP-based glycine sensor)[4] | GGCCCGGAUAGCUCAGUCGGUAGAGCAGCGGCCGG UUGCCAUGUGUAUGUGGGCCAAUCGUGGCGUGUCG GCCUGCUUCGGCAGGCACUGGCGCCGCCCACAUACU CUGAUGAUCCUGCCGAAUGAAGUUUAAUAGAAGUG GGGAAUCGUUUGAUUUUCCAUGACUGUAAAUGGAC GGAACUCUGGAGAGACCGUAAAGGCACCGAAGGGG CAAGGCCUGUCGAGUAGAGUGUGGGCUCGCAAGAG ACGGUCGGGUCCAGGCUCAAACUCUCAGGUAAAAG GACAGAGUUGGAUCAUUCAUGGCAACGGCCGCGGG UCCAGGGUUCAAGUCCCUGUUCGGGCGCCA |
| | .(((((((..((((........)))).((((((.(((((((((.((((((((((.......(((((....((((((....))) )))....)))))))))))))))...((((((((.. (((......[((((.[((..[(((([[((((....))))]])))]..))]))))]...)))..(((((......((.....)).( ((..((([(....[((((((..((...((.[((...(((....)))))].))...)))))))]....)]))..))).......)))) ) ..)))))))))))))))))))))))).....(((((.......)))))))))))).... |
| Test case 5 (MgZ)[5] | GATGTGTCCGTGCTATGGTTCGATTCTTGATCAACCA GGTCGGGGCCGAAATATAGGATATTTTGGGAGGCTA TGCTAGGAAG |
| | ......(((..((..(((........(((..))))))........(([(((((((...)))))))]))........))..)))).. |

| | |
|---|---|
| Test case 6 (NCB)-1[6] | CCCCCTTAATCCCCCTATAATAAATTTTAAATATTATT TATTAAT |
| | ............................................. |
| Test case 6 (NCB)-2[6] | ATTAATAAATAATATTTAAAATTTATTATAGGGTGGG GTGGGGTGGGG |
| | ................................................ |
| Test case 7 (strand-displacement aptasensor)-F[7] | CTCTCTCGCTTGACTTGTTTGTAATGTAGGGTTCCGAG CGA |
| | ......................................... before displacement |
| | .....(((((([(....................)])))))) after displacement |
| Test case 7 (strand-displacement aptasensor)-Q[7] | AAGCGAGAGAG |
| | ........... |
| Test case 8 (aptamer switch)[8] | CACCTGGGGGAGTATTGCGGAGGAAGGTTTTTTTTT TTTTTTCCCCCAGGTG |
| | (((((((......................................))))))) quenched |
| | .................................................... unfolded |
| | ((((...((.......))...)))).......................... target-bound |

**Table S2. Two-strand complex duplex regions**

| Complex name | Strands | Duplex region(s) |
|---|---|---|
| Main text **Fig. 1** bottom | Main text **Fig. 1** bottom-1 | 10-13 |
| | Main text **Fig. 1** bottom-2 | 1-4 |
| Main text **Fig.2** | Main text **Fig.2**-Strand A | 15-19 |
| | Main text **Fig.2**-Strand B | 1-5 |
| Test case 3 (internal loop)[3] | Test case 3 (internal loop)-1 | 1-4, 10-13 |
| | Test case 3 (internal loop)-2 | 5-8, 1-4 |
| Test case 6 (NCB)[6] | Test case 6 (NCB)-1 | 16-45 |
| | Test case 6 (NCB)-2 | 1-30 |
| Test case 7-1 (strand-displacement aptasensor)[7] | Test case 7 (strand-displacement aptasensor)-F | 1-11 |
| | Test case 7 (strand-displacement aptasensor)-Q | 1-11 |

**Supplementary Figure 1**

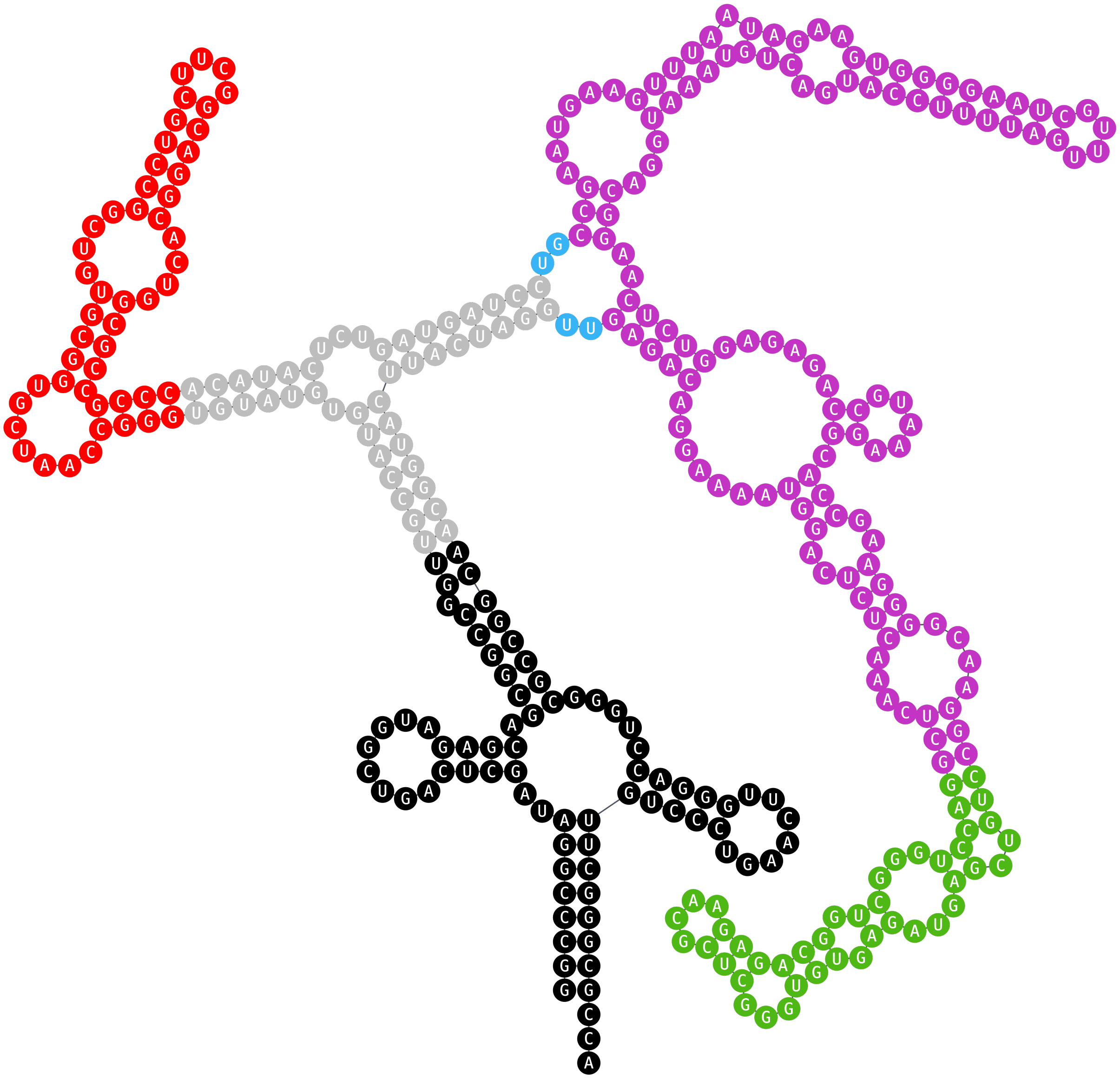

**Fig. S1. Test case 4 (R-FAP-based glycine sensor) with nucleotide labels.** The AptaRender reconstruction demonstrates its ability to visualize complex structures in greater detail than a simplified bond-line representation.

**Supplementary Figure 2**

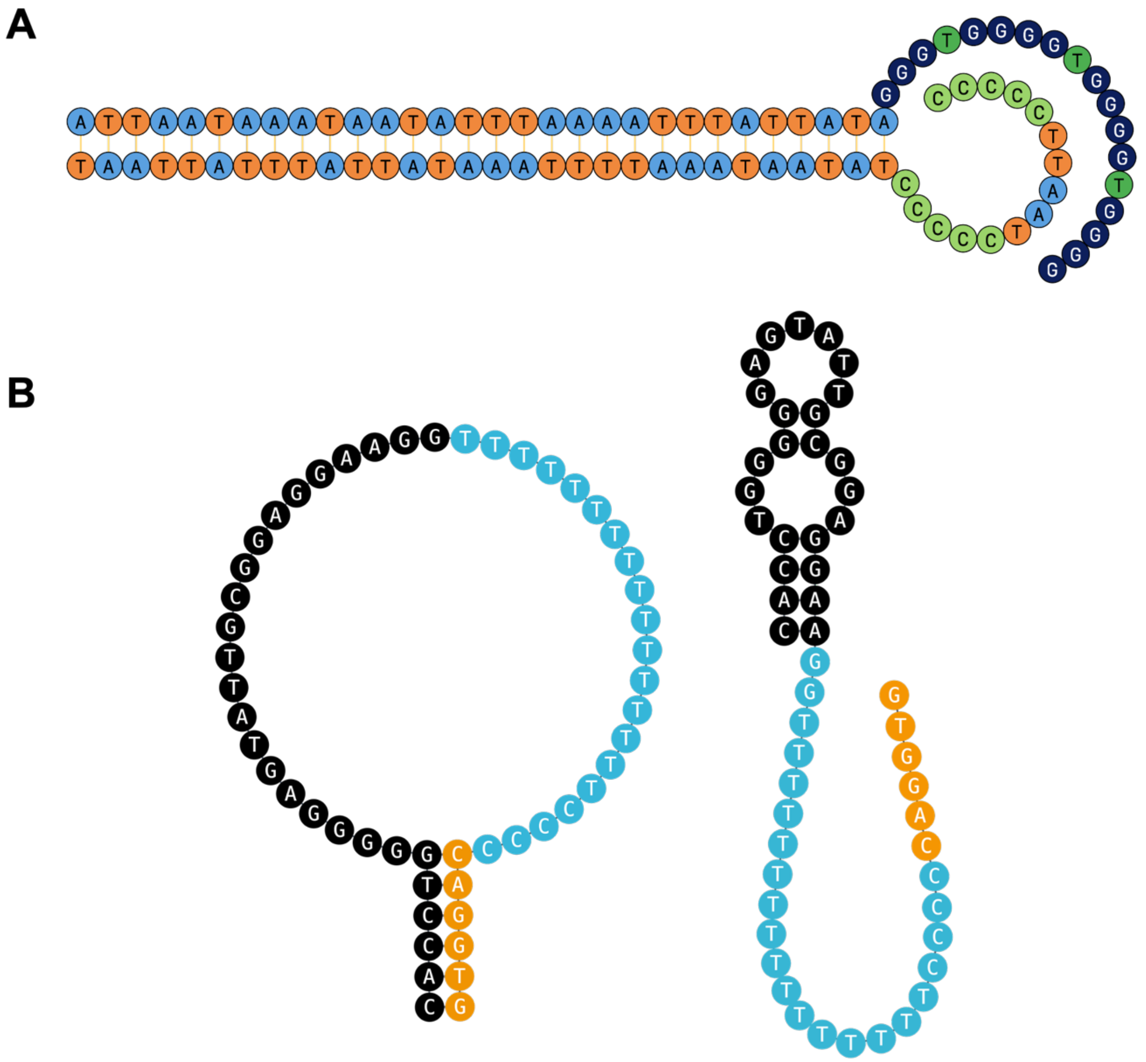

**Fig. S2. Test case 6 (NCB) and Test case 8 (aptamer switch) with nucleotide labels.** Together with Fig. S1, these three examples showed that AptaRender reconstruction demonstrates its ability to visualize complex structures in greater detail than a simplified bond-line representation.

**Files S1-S32 (Separate files)**

**AptaRender-reproduced tests cases exported as PNG (1139×3304 px) and SVG files**
All structures reproduced using AptaRender were exported in both PNG and SVG formats which were also presented in the manuscript (Main text **Fig. 3B, D, F, H, J, L, N, P,** Supplementary **Fig. S1-2**).

**User tutorial (Separate files)**

**Comprehensive user tutorial for AptaRender 2.2.4**, as a Microsfot Word file.

### AptaRender

#### User tutorial: every tab, button, gesture and shortcut

AptaRender turns a sequence and a dot-bracket structure you supply into an editable 2D figure of a nucleic-acid secondary structure, ready to export as SVG or PNG for papers, posters and slides. It does **not** predict structures. You tell it the pairing, and it gives you full control over how the result looks.

This guide walks through the screen, a five-minute quick start, each tab of the control panel, everything you can do with the mouse on the canvas, the keyboard shortcuts, and a few step-by-step recipes for common figures.

*Covers the build of October 2026, including box select, parts, rotate selection and zoom to selection.*

| Section | Topic |
| --- | --- |
| 1 | The screen at a glance |
| 2 | Quick start |
| 3 | Writing a structure |
| 4 | The control panel, tab by tab |
| 5 | Working on the canvas (mouse) |
| 6 | Keyboard shortcuts cheat sheet |
| 7 | Recipes for common figures |
| 8 | Saving, undo, and where your data lives |
| 9 | Limitations and troubleshooting |

#### References

(1) Kim, S.; Hong, S.; Walker, J. N.; He, Y.; Nguyen, A.-T.; Chen, W.-R.; Seifi, S.; Chen, Y.-I.; Kuo, Y.-A.; Brodbelt, J. S.; Yeh, H.-C. Mechanistic Insights into the Color Transformation of a Non-FRET Substrate for RNase Activity Detection. bioRxiv March 13, 2026, p 2026.03.11.710959. https://doi.org/10.64898/2026.03.11.710959

(2) Kuo, Y.-A.; Chen, Y.-I.; Siraj, N.; He, Y.; Yang, Z.; Wang, Y.; Batchelder-Schwab, E. J.; Korkmaz, Z.; Yonas, S.; Nguyen, T. D.; Hong, S.; Nguyen, A.-T.; Kim, S.; Seifi, S.; Fan, P.-H.; Wu, Y.; Liu, H.-W.; Lu, Y.; Ren, P.; Mao, C.; Yeh, H.-C. Fluorogenic Aptamer Optimization on a Massively Parallel Sequencing Platform. *ACS Sens.* **2026**, *11* (5), 3587–3600. https://doi.org/10.1021/acssensors.5c04046

(3) Monferrer, A.; Zhang, D.; Lushnikov, A. J.; Hermann, T. Versatile Kit of Robust Nanoshapes Self-Assembling from RNA and DNA Modules. *Nat Commun* **2019**, *10* (1), 608. https://doi.org/10.1038/s41467-019-08521-6

(4) Bodin, M. R.; Hammond, M. C. Visualizing Intracellular Glycine with Two-Dye and Single-Dye Ratiometric RNA-Based Sensors. *Nucleic Acids Res* **2025**, *53* (17), gkaf839. https://doi.org/10.1093/nar/gkaf839

(5) Chiuman, W.; Li, Y. Simple Fluorescent Sensors Engineered with Catalytic DNA 'MgZ' Based on a Non-Classic Allosteric Design. *PLOS ONE* **2007**, *2* (11), e1224. https://doi.org/10.1371/journal.pone.0001224

(6) Kuo, Y.-A.; Jung, C.; Chen, Y.-A.; Kuo, H.-C.; Zhao, O. S.; Nguyen, T. D.; Rybarski, J. R.; Hong, S.; Chen, Y.-I.; Wylie, D. C.; Hawkins, J. A.; Walker, J. N.; Shields, S. W. J.; Brodbelt, J. S.; Petty, J. T.; Finkelstein, I. J.; Yeh, H.-C. Massively Parallel Selection of NanoCluster Beacons. *Advanced Materials* **2022**, *34* (41), 2204957. https://doi.org/10.1002/adma.202204957

(7) He, Y.; Yang, Z.; Kuo, Y.-A.; Wu, Y.; Fonseca-Albert, D.; Le, K. K.; Guo, J.; Wang, Y.; Nguyen, A.-T.; Chen, Y.-I.; Kim, S.; Chen, W.-R.; Seifi, S.; Hong, S.; Nguyen, T. D.; Chen, Y.; Ren, P.; Lu, Y.; Yeh, H.-C. High-Throughput On-Chip Screening Enables Rapid Adaptation of DNA Aptamers to SARS-CoV-2 Evolution. *ACS Nano* **2026**, *20* (34), 23941–23957. https://doi.org/10.1021/acsnano.6c08330

(8) Wilson, B. D.; Hariri, A. A.; Thompson, I. A. P.; Eisenstein, M.; Soh, H. T. Independent Control of the Thermodynamic and Kinetic Properties of Aptamer Switches. *Nat Commun* **2019**, *10* (1), 5079. https://doi.org/10.1038/s41467-019-13137-x
