## Supplementary figures and images for "AptaRender: an interactive drawing and styling tool for 2D nucleic acid structures"

### case1.png

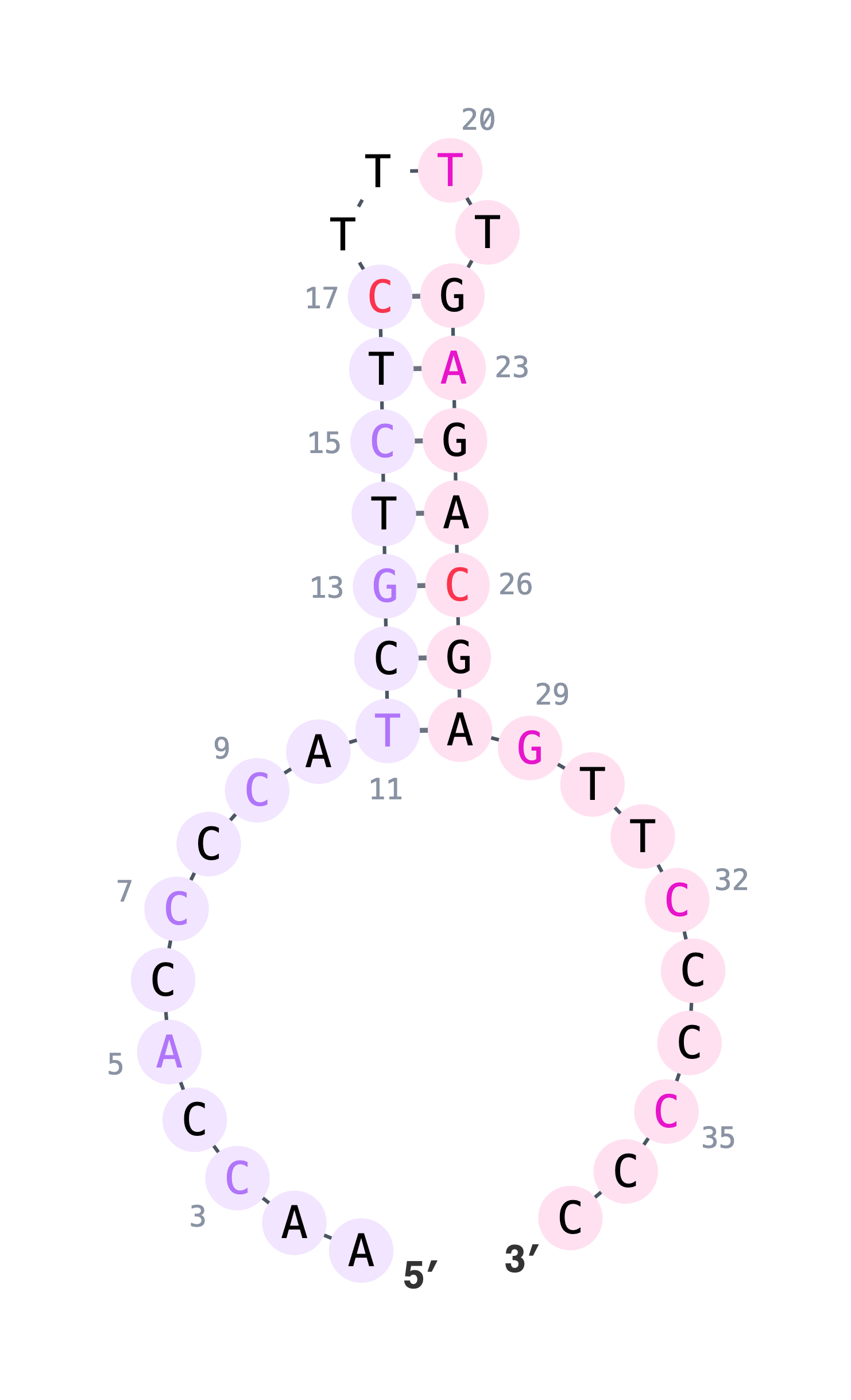

### case2.png

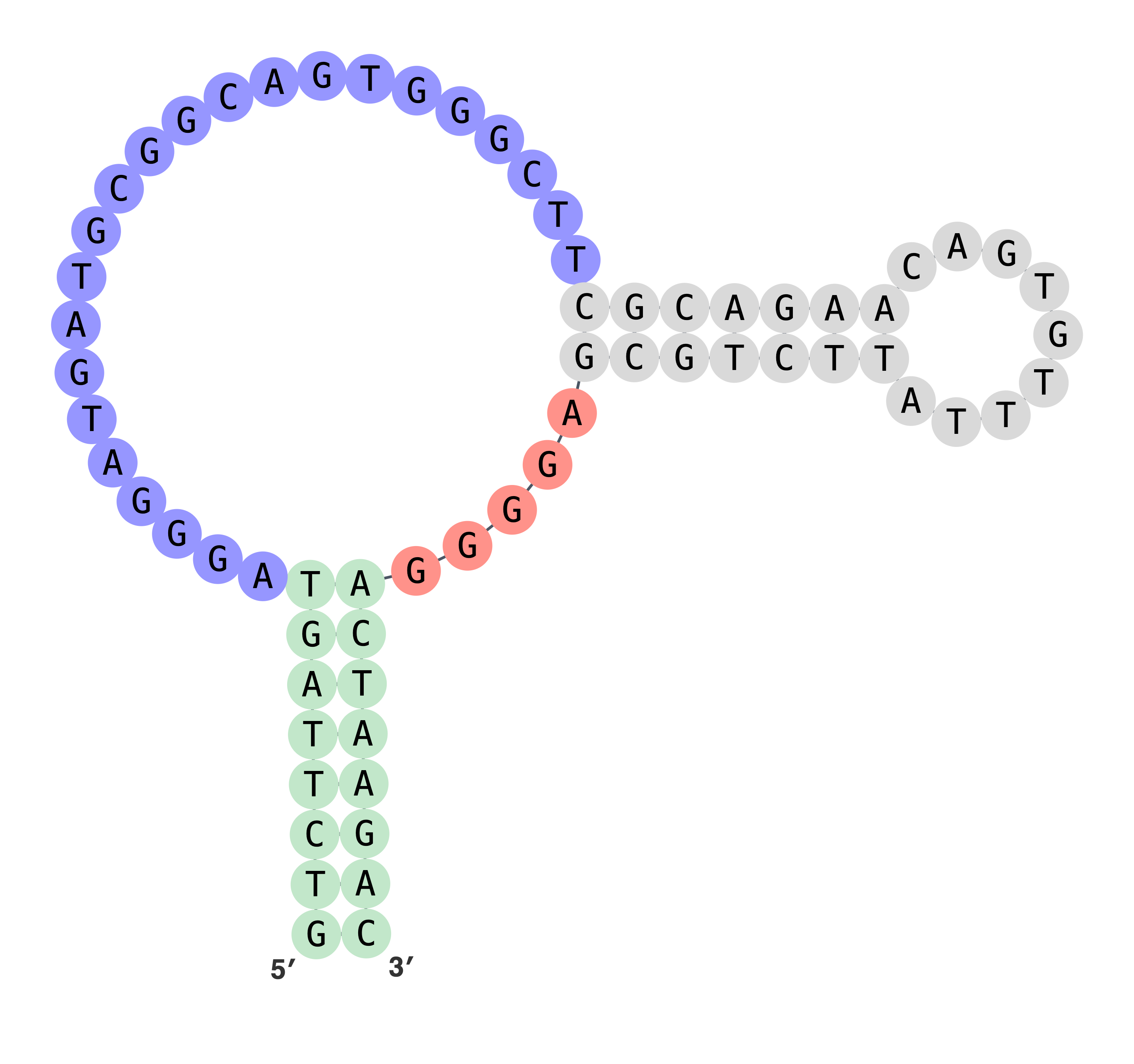

### case3.png

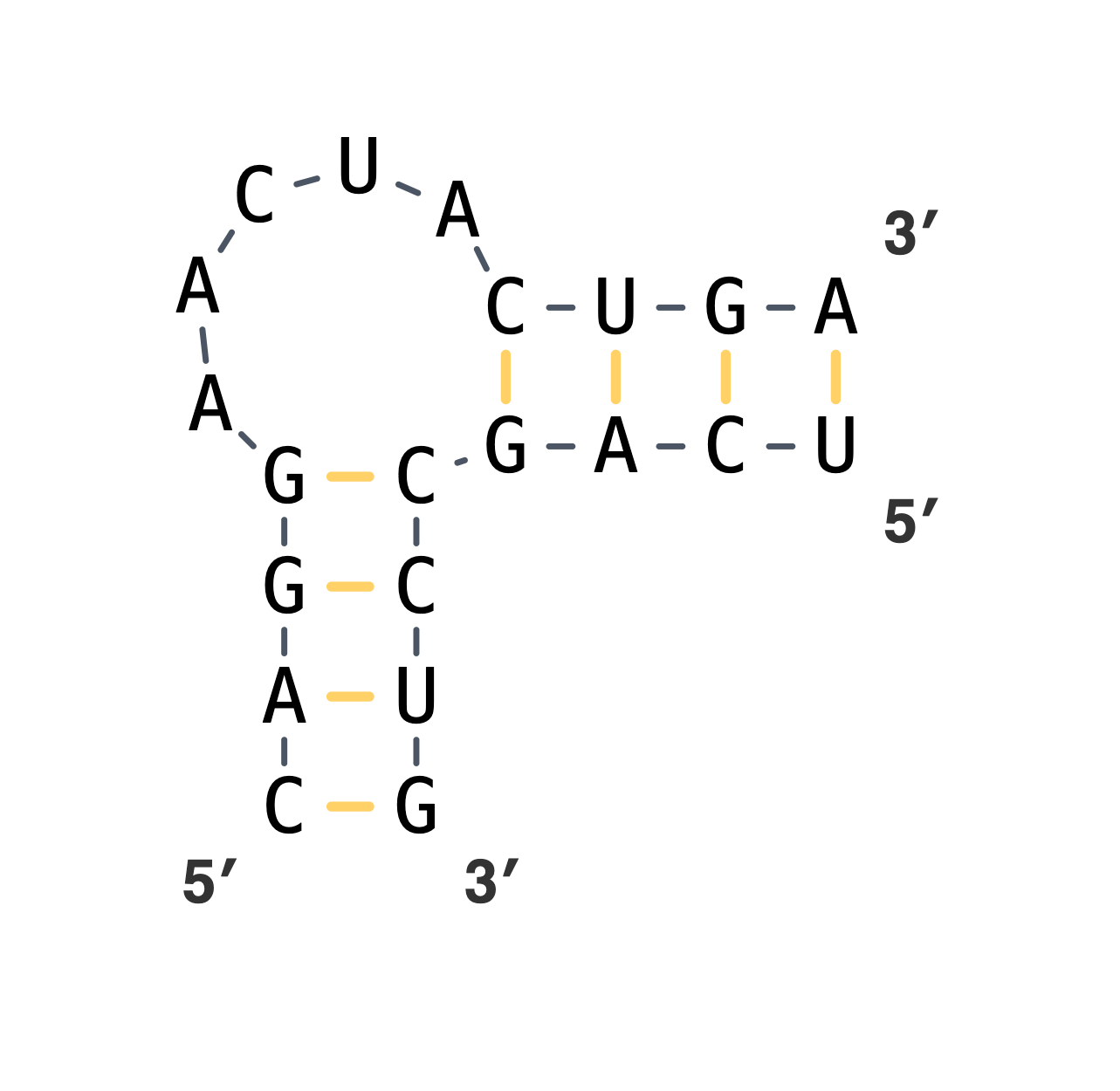

### case4.png

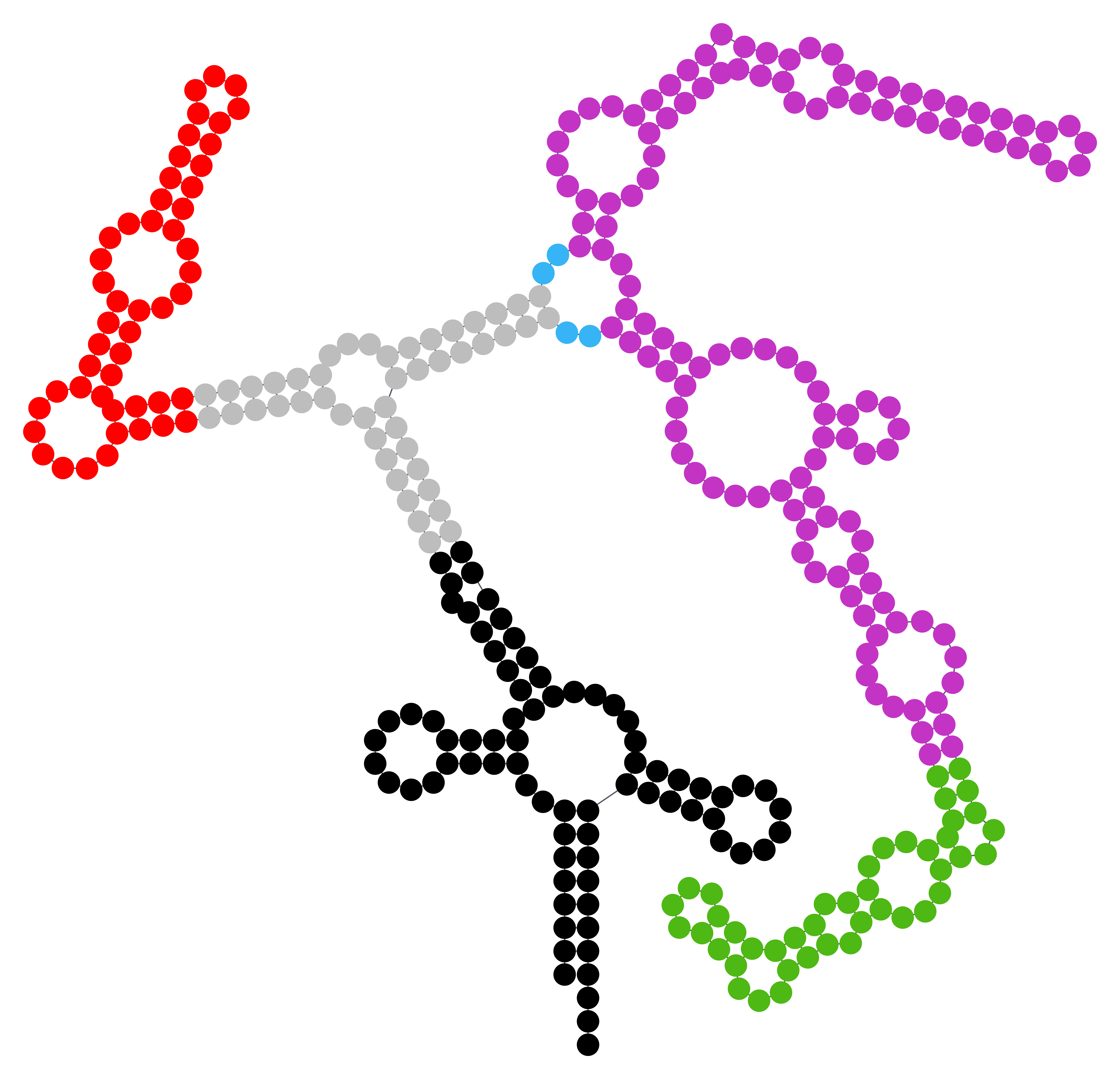

### case4_with nt labels.png

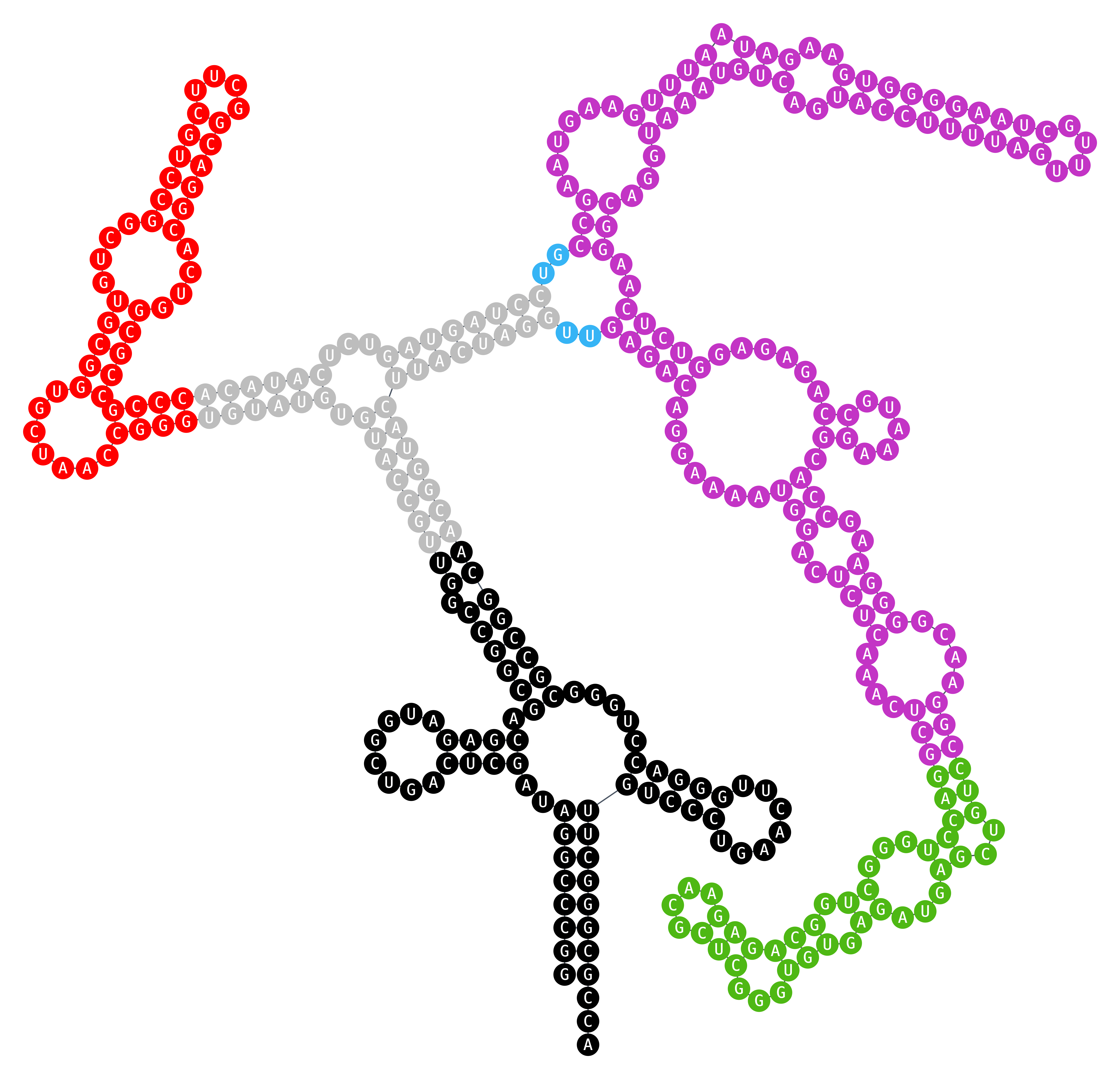

### case5.png

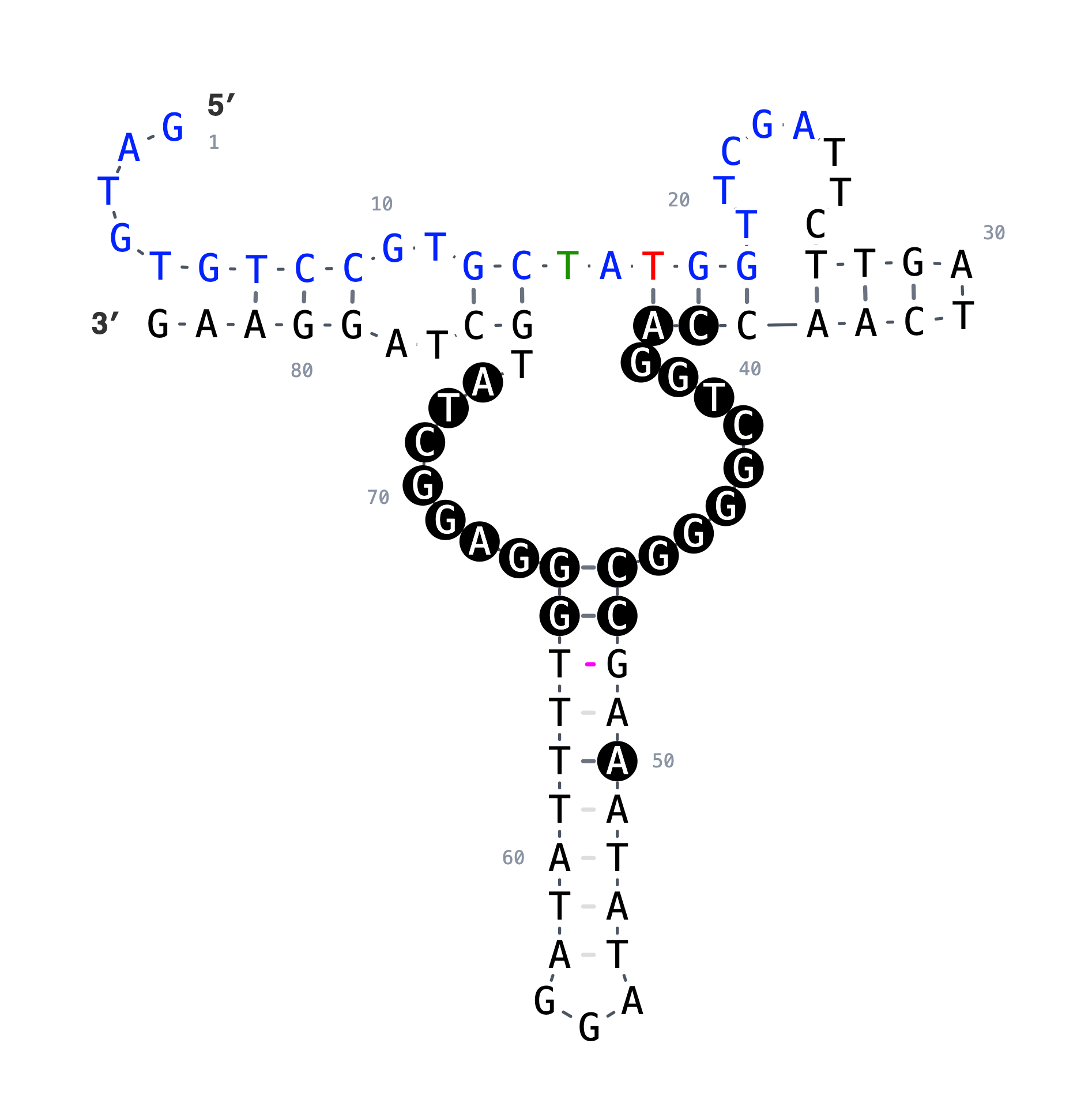

### case6.png

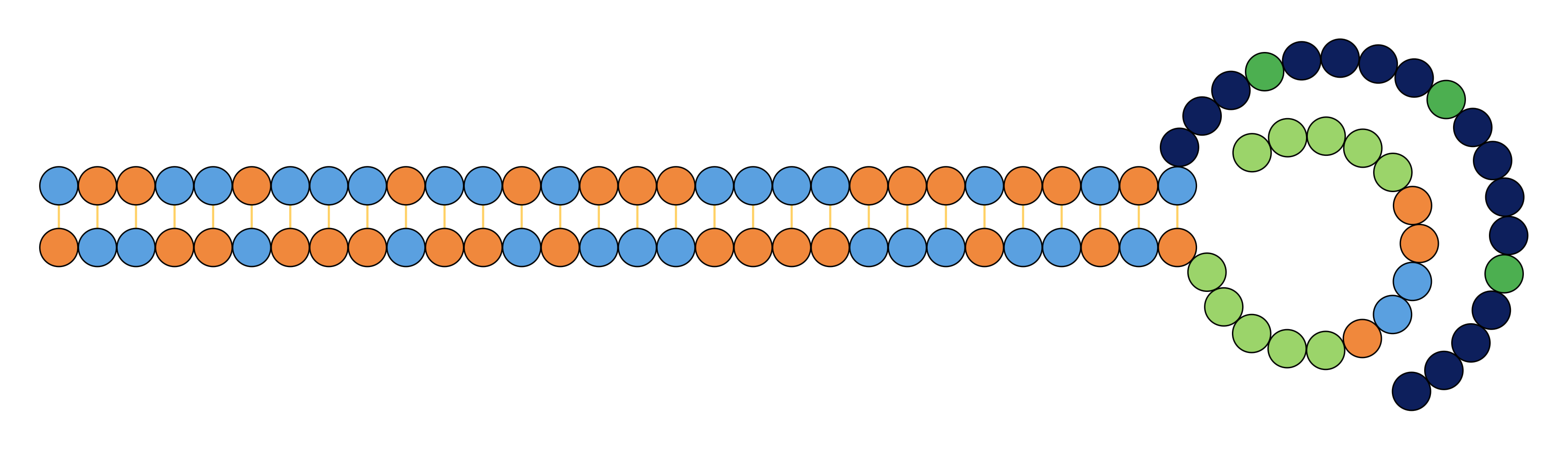

### case6_with nt labels.png

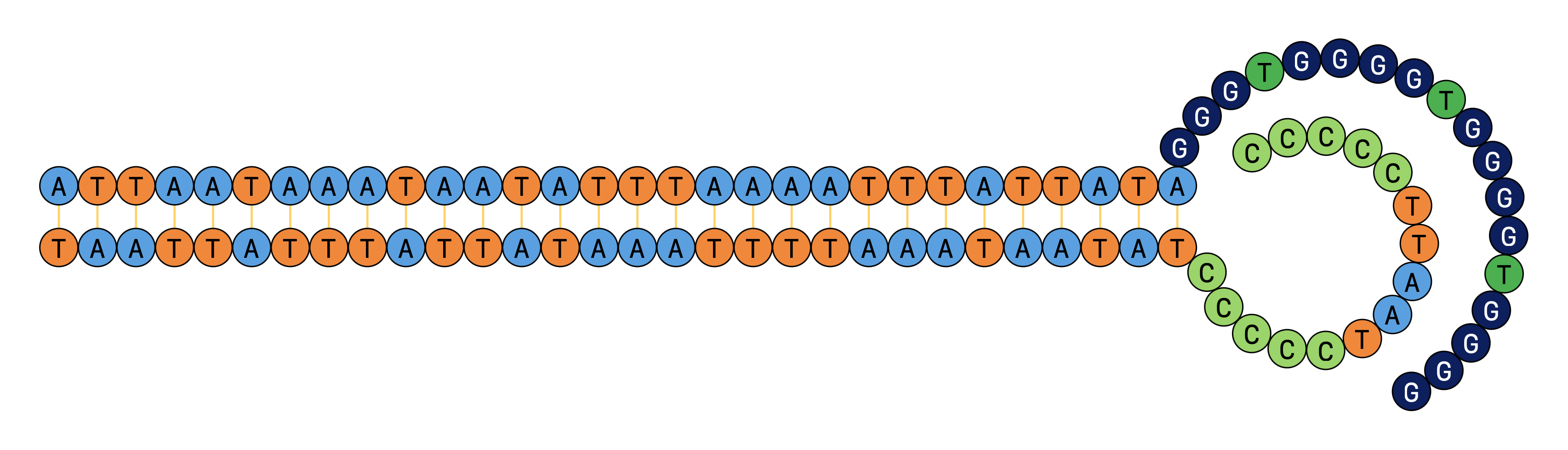

### case7-1.png

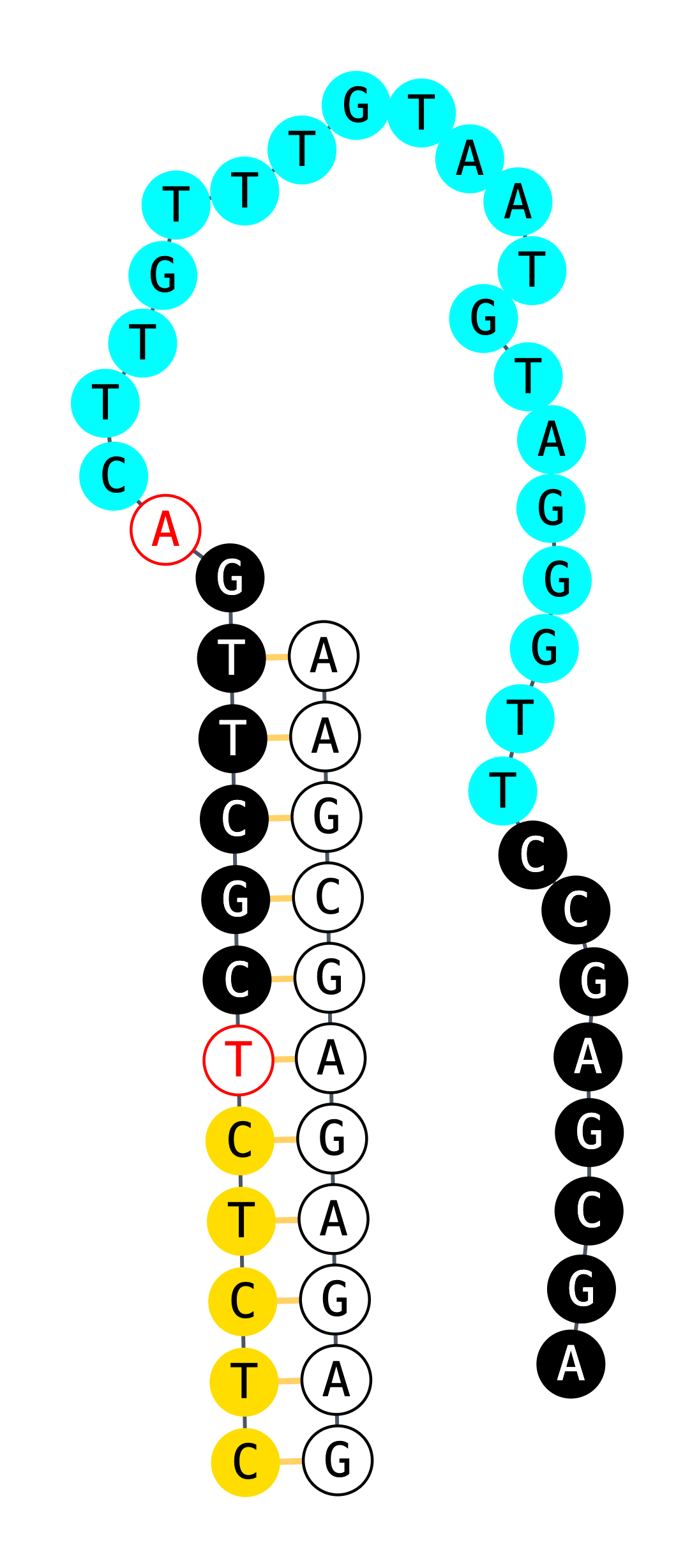

### case7-2.png

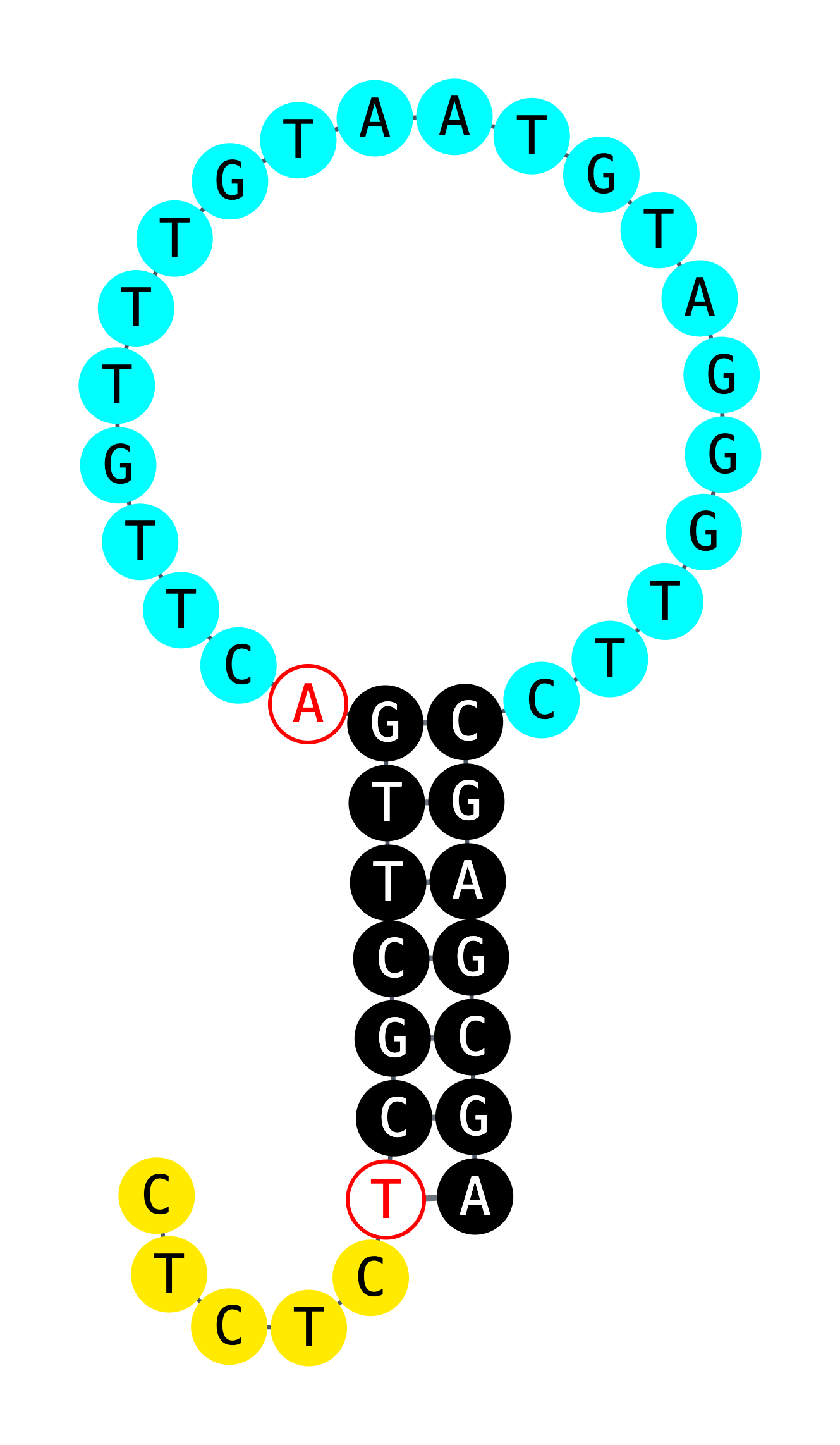

### case7-3.png

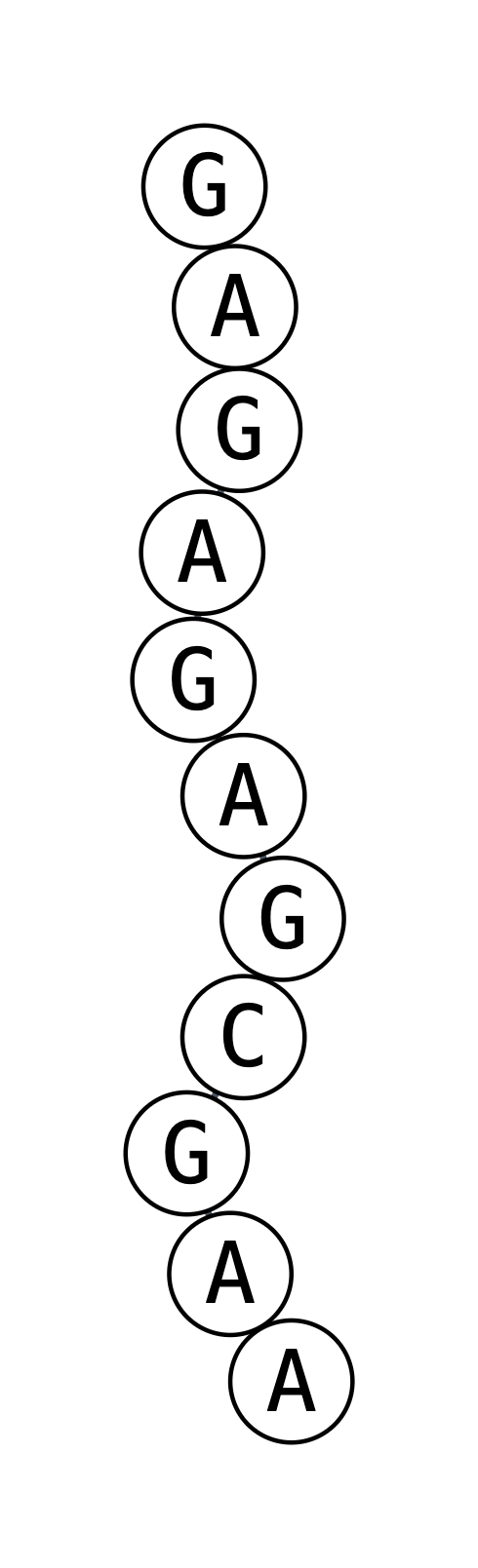

### case8-1.png

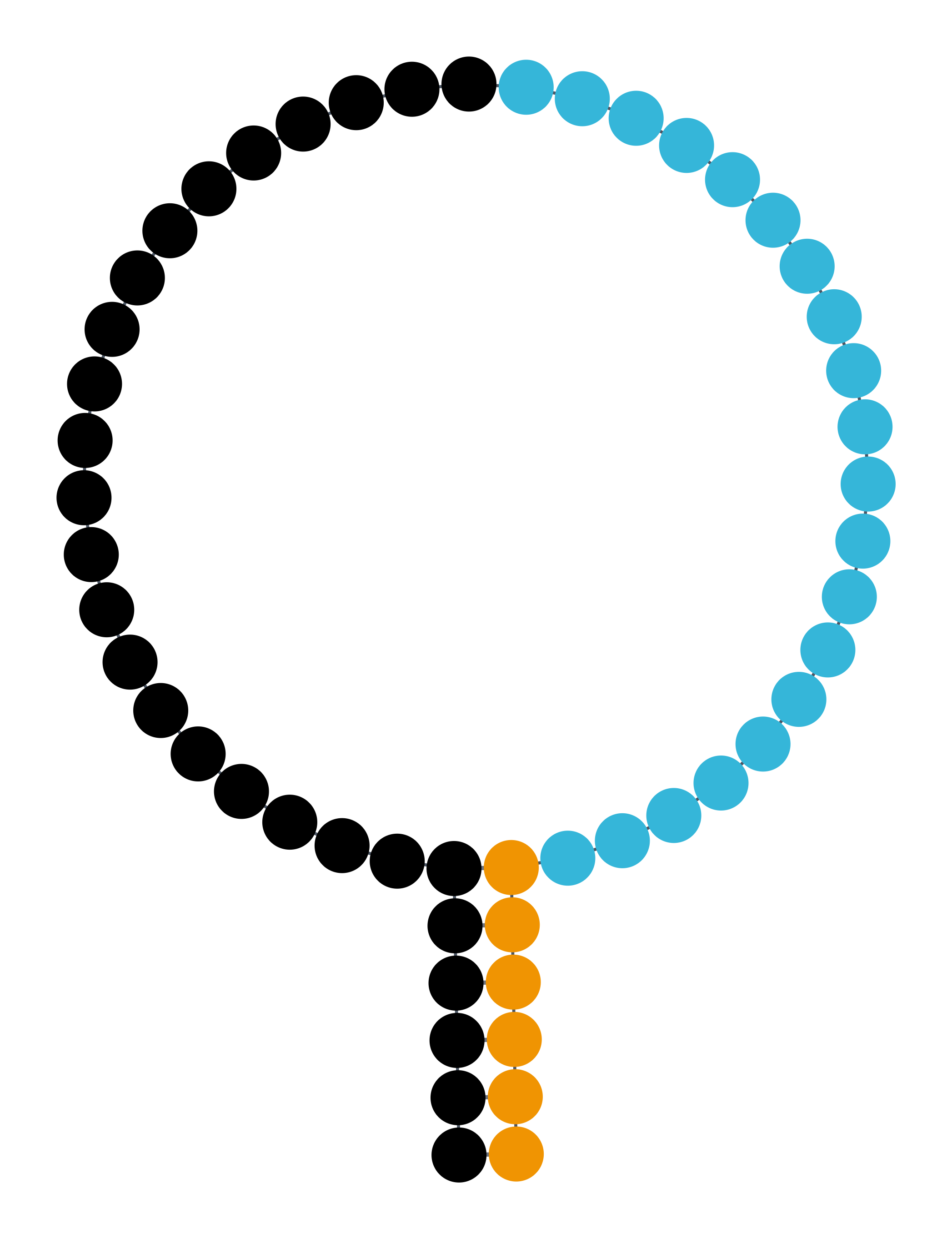

### case8-1_with letter.png

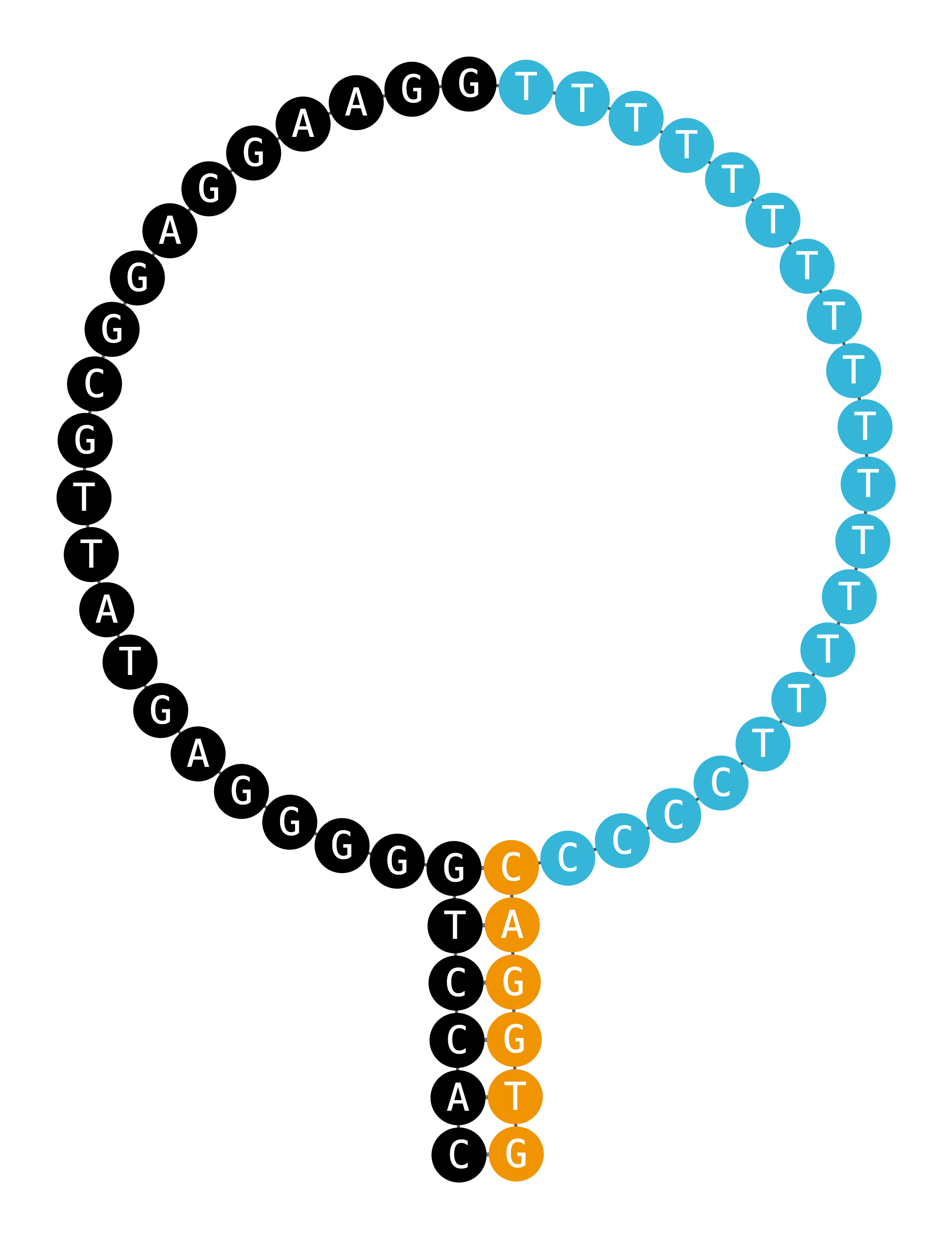

### case8-2.png

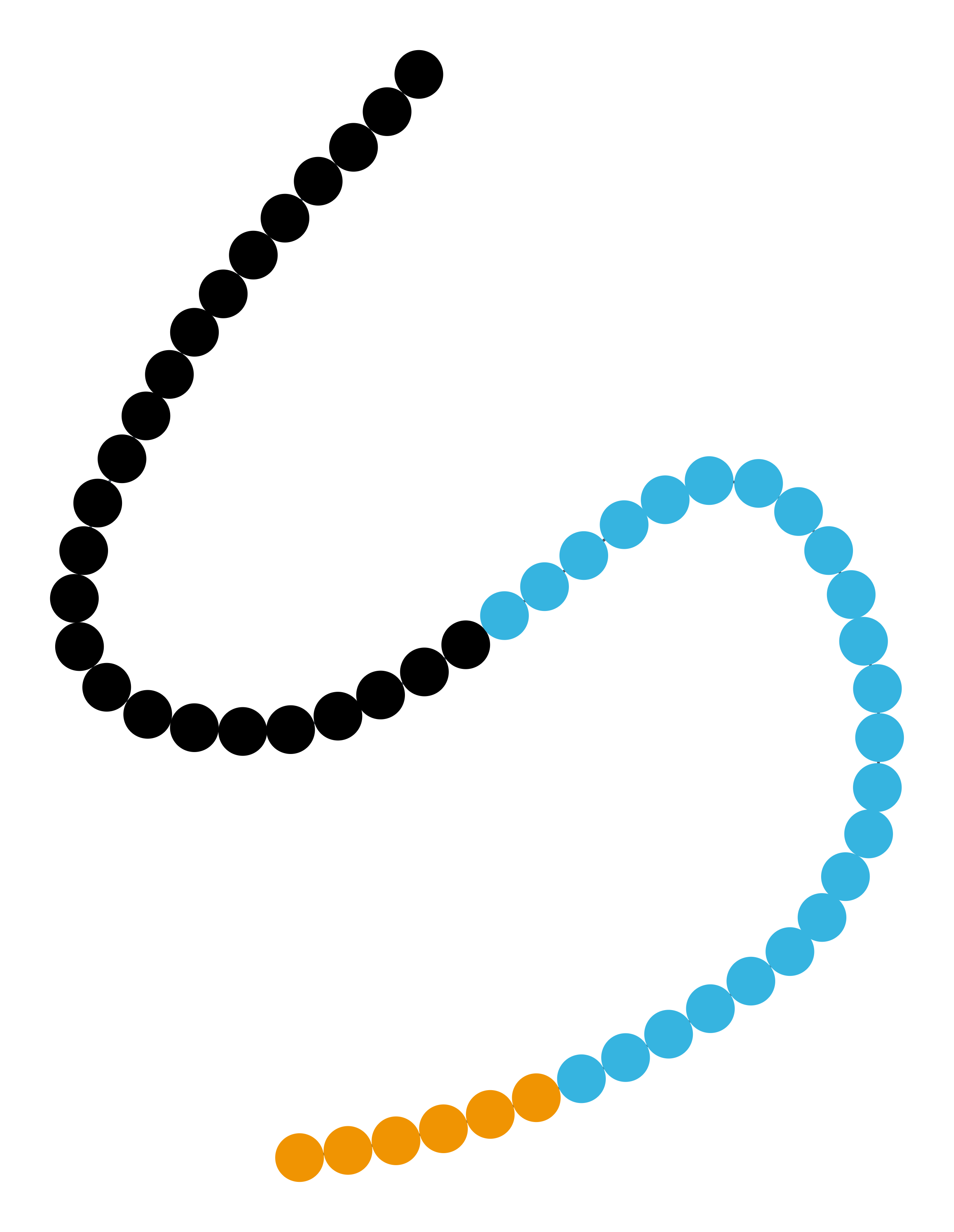

### case8-3.png

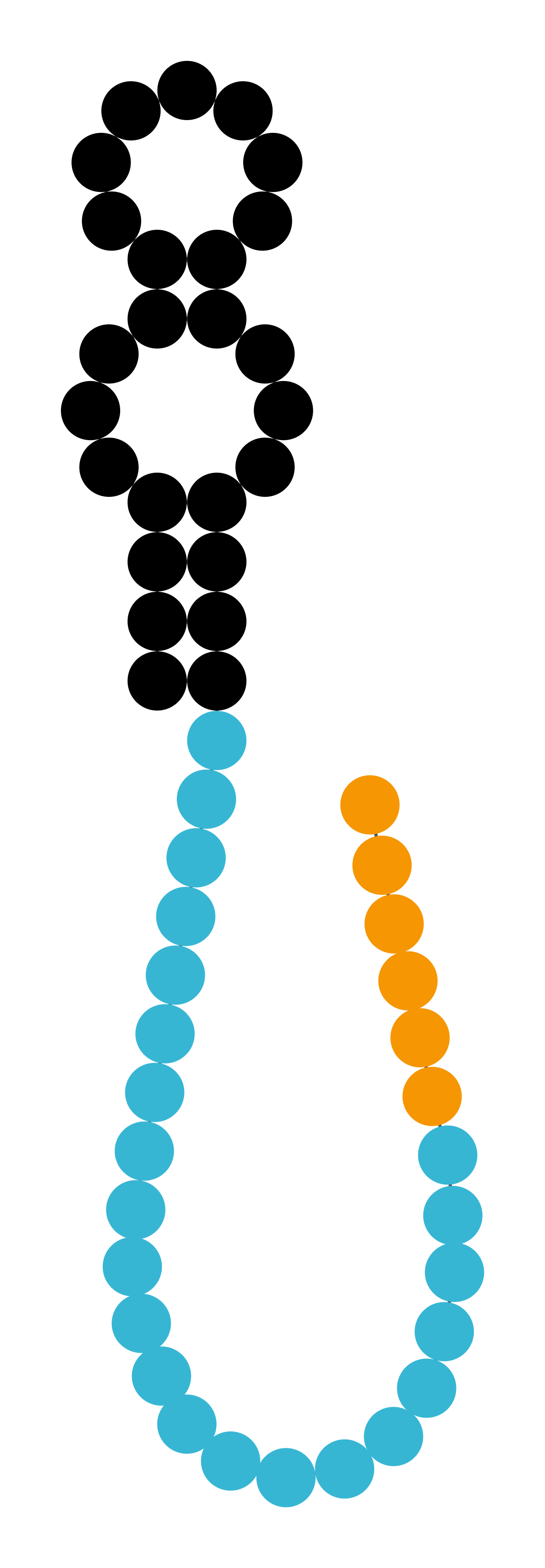

### case8-3_with letter.png

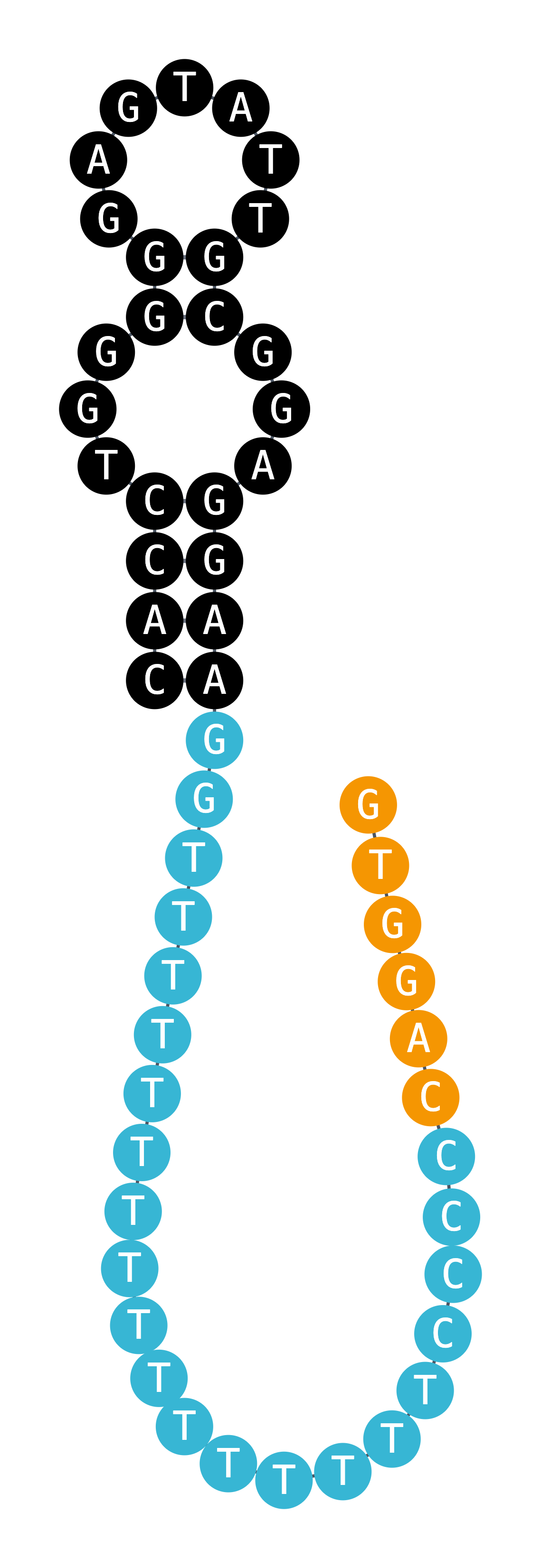
