## Supplementary material for "AptaRender: an interactive drawing and styling tool for 2D nucleic acid structures": User tutorial

**AptaRender**

User tutorial: every tab, button, gesture and shortcut

AptaRender turns a sequence and a dot-bracket structure you supply into an editable 2D figure of a nucleic-acid secondary structure, ready to export as SVG or PNG for papers, posters and slides. It does **not** predict structures. You tell it the pairing, and it gives you full control over how the result looks.

This guide walks through the screen, a five-minute quick start, each tab of the control panel, everything you can do with the mouse on the canvas, the keyboard shortcuts, and a few step-by-step recipes for common figures.

*Updated November 2026 for the Bases tab (letters and circles together), separate letter and circle sizes, and the (i) help buttons.*

| **Section** | **Topic** |
| --- | --- |
| 1 | The screen at a glance |
| 2 | Quick start |
| 3 | Writing a structure |
| 4 | The control panel, tab by tab |
| 5 | Working on the canvas (mouse) |
| 6 | Keyboard shortcuts cheat sheet |
| 7 | Recipes for common figures |
| 8 | Saving, undo, and where your data lives |
| 9 | Limitations and troubleshooting |

### 1. The screen at a glance

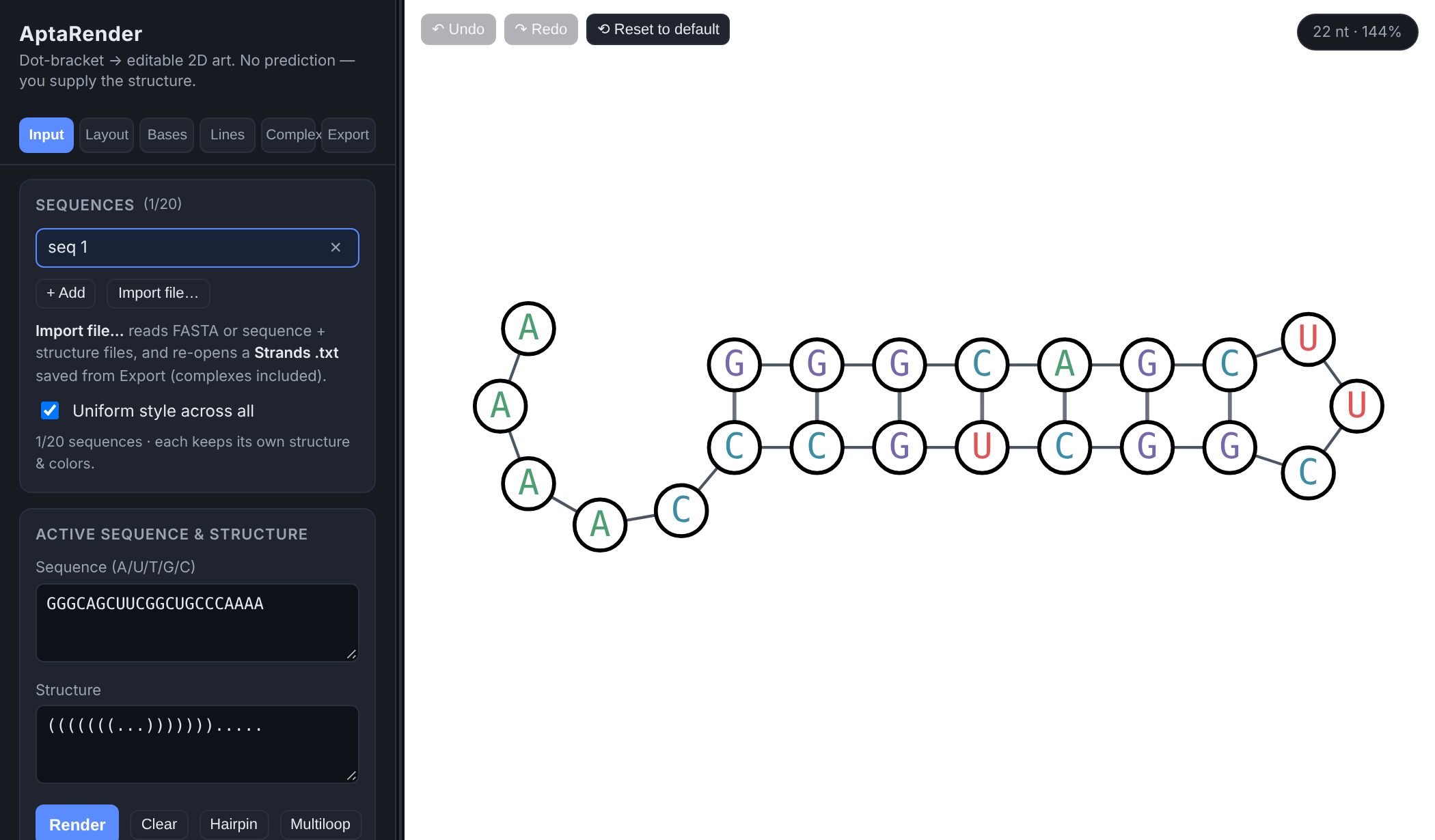

*Figure 1. The AptaRender window with the built-in Hairpin example loaded.*

| **Area** | **What it is** |
| --- | --- |
| **Control panel** (left) | All settings, grouped into six tabs along the top: Input, Layout, Bases, Lines, Complex, Export. A box with an **(i)** at the top right of its heading has a short how-to behind it (see section 4). |
| **Selection bar** | Appears under the tabs on Layout, Bases and Complex. Shows how many bases are selected, with **Select all**, **Clear** and **Zoom** buttons (see 4.0). |
| **Canvas** (right) | The drawing. Pan, zoom, select and drag bases here (see section 5). |
| **Undo / Redo** (canvas, top-left) | Step backward or forward through your edits. Greyed out when there's nothing to undo or redo. |
| **Reset to default** (next to Undo/Redo) | Puts what's open back to a fresh render: manual positions, letter colors (back to the palette), circle fills, bond colors, base sizes and rotation are cleared, and every style setting (letter and circle sizes, circles, lines, numbering, tail style, background) returns to its default. With **Uniform style** on, the style part applies to all sequences. One **Undo** brings everything back. |
| **Badge** (canvas, top-right) | Length of what's on screen and the current zoom, e.g. "22 nt · 206%". In a complex it reads "13+13 nt (complex)". |
| **Divider** | The thin strip between panel and canvas. Drag it to make the panel wider or narrower; the width is remembered. |

### 2. Quick start

1. On the **Input** tab, paste your sequence into **Sequence** (A, U, T, G, C or N; spaces and line breaks are ignored, lowercase is fine).
2. Paste the matching dot-bracket string into **Dot-bracket structure**. It must be exactly the same length as the sequence.
3. Click **Render**. The structure is drawn and fitted to the canvas. If something's wrong, a red message under the buttons says exactly what and where.
4. Use **Bases** (letters and circles) and **Lines** to style it, and **Layout** to rotate or rearrange it. Drag bases on the canvas to fine-tune positions.
5. On **Export**, pick a PNG scale (or go straight to SVG) and click **SVG** or **PNG** to download the figure.

**Tip** Click **Hairpin** or **Multiloop** on the Input tab to load a ready-made example into the current sequence slot. It's the fastest way to try every feature in this guide.

### 3. Writing a structure

The structure is one character per base, aligned with the sequence. Pairs must nest inside each other (no crossing pairs).

| **Character** | **Meaning** |
| --- | --- |
| .  (also  -   :   , ) | Unpaired base. |
| (  …  ) | A Watson-Crick base pair, drawn as a **solid** rung. |
| [  …  ]   {  …  }   <  …  > | A non-Watson-Crick pair, drawn as a **dashed** rung. Use these to mark wobble, Hoogsteen or other non-canonical pairs so they stand out. |

Example:  GGGCAGCUUCGGCUGCCCAAAA  with  (((((((...))))))).....  is a 7-bp hairpin with a 3-nt loop and a 5-nt 3′ tail. Changing the middle of the stem to  ((([[[(...)]]]))).....  would draw those three pairs dashed.

**Tip** Copying from PowerPoint, Word, PDFs or web pages is fine. Hidden characters they add (zero-width spaces, soft hyphens, non-breaking spaces, line separators) are removed automatically, an autocorrected "…" is read as three dots, and full-width brackets are read as normal ones.

##### Error messages you might see

| **Message** | **How to fix it** |
| --- | --- |
| Length mismatch | The sequence and structure have different lengths. The message gives both numbers. |
| Unexpected character … in sequence | Only A U T G C N are allowed in the sequence. The message gives the position, and names the character by its code (e.g. U+00A9) if it's hard to see. |
| Unbalanced "(" / ")" at position N | An opening bracket was never closed, or a closing bracket has no partner. Count the brackets around position N. |
| Mismatched bracket | A pair was opened with one bracket type and closed with another, e.g.  [  closed by  ) . Use the same type on both ends. |

### 4. The control panel, tab by tab

**Tip** Many boxes have a small **(i)** at the right of their heading. Click it to read a short how-to for that box; click anywhere else, press Esc or switch tabs to close it. The Input, Complex and Export tabs, and the color-file format, keep their instructions on screen.

#### 4.0 The selection bar

Most styling and layout actions work on the **selected** bases (orange rings on the canvas). The selection bar is your shortcut to manage it:

| **Control** | **What it does** |
| --- | --- |
| **n selected** | How many bases are currently selected (across both strands in a complex). |
| **Select all** | Selects every base in the current view. |
| **Clear** | Deselects everything. Clicking empty canvas does the same. |
| **Zoom** | Centers the canvas on the selected bases and zooms in to fit them. With nothing selected it behaves like **Fit to view**. |

#### 4.1 Input tab: sequences and structure

##### Sequences list

| **Control** | **What it does** |
| --- | --- |
| Sequence name | Click to open that sequence. **Double-click** to rename it. Empty slots are tagged "· empty". |
| **×** next to a name | Deletes that sequence. Any complex built from it is deleted too (a note tells you how many). |
| **+ Add** | Adds a new, blank sequence slot and opens it. Up to **20** sequences. |
| **Import file…** | Loads one or more sequences from a FASTA-style file (.fa, .fasta, .txt, .dbn, .seq, .dot). Each record is a  >name  line, then the sequence line(s), then optionally the dot-bracket line(s). Extras beyond the 20-sequence limit are skipped. |
| **Uniform style across all** | On (default): every sequence shares one look (letter and circle sizes, colors, circles, lines, numbering, tail style, background). Off: each sequence remembers its own look. Per-base colors and dragged positions always belong to each sequence either way. |

##### Active sequence and structure

| **Control** | **What it does** |
| --- | --- |
| **Sequence** box | The sequence for the open slot. Saved as you type. Long sequences wrap onto the next line. |
| **Structure** box | The dot-bracket structure for the open slot (see section 3). It wraps at the same column as the Sequence box, so each structure character sits under its base. |
| **Render** | Draws the structure from the two boxes, sets the rotation to 0° and fits it to the view. |
| **Clear** | Empties the open slot's sequence, structure and colors. The slot stays in the list. |
| **Hairpin** / **Multiloop** | Loads an example into the open slot, replacing what's there, and renders it. |

**Watch out Render** starts that sequence fresh: custom letter and circle colors, bond colors, base sizes, dragged positions and rotation are reset. If you only meant to restyle, use the other tabs instead. If you clicked it by accident, press  Ctrl + Z .

#### 4.2 Layout tab: orientation and positions

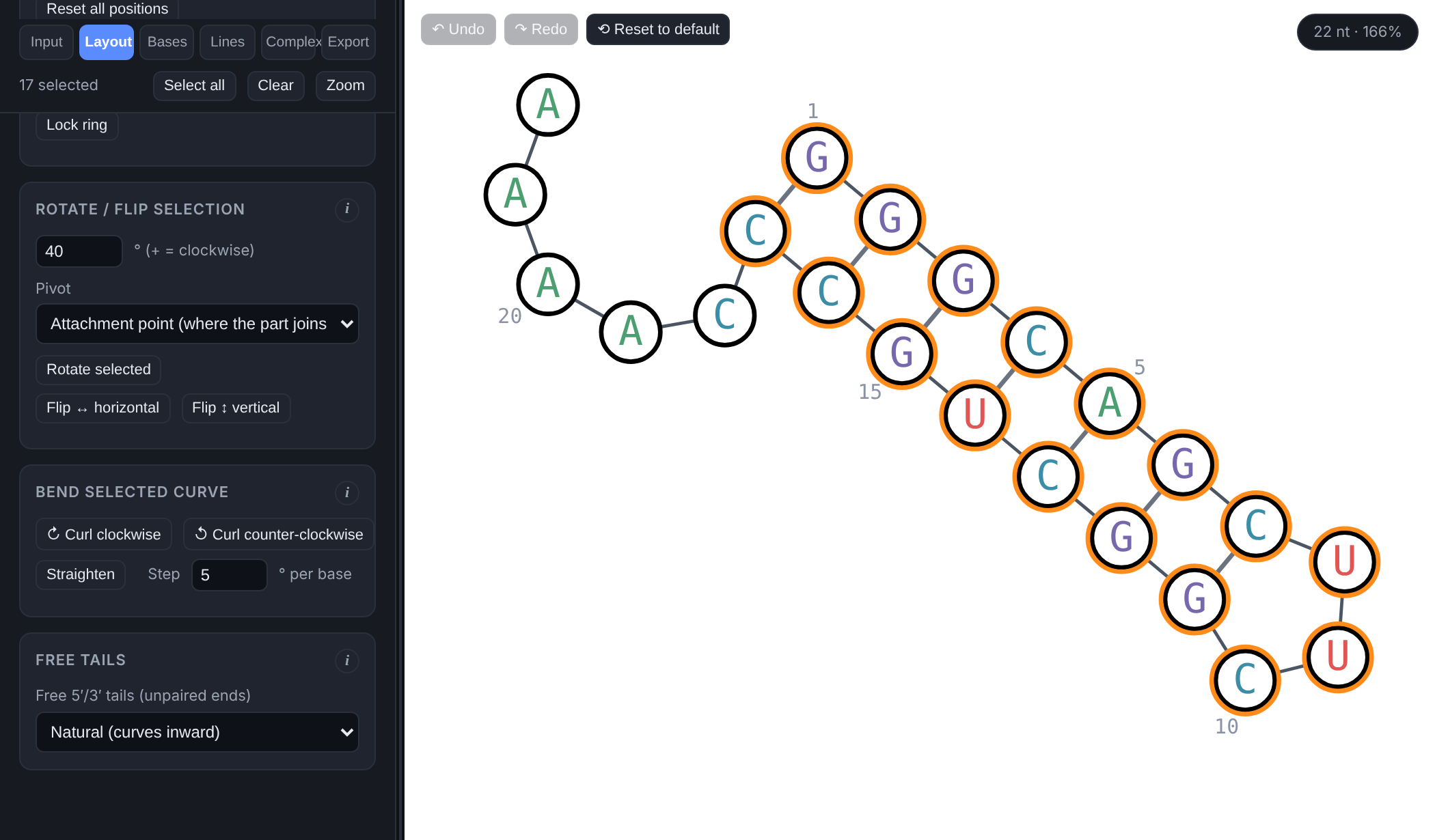

*Figure 2. Layout tab. The hairpin arm was selected and rotated 40° about its base pair; the tail stayed put.*

| **Control** | **What it does** |
| --- | --- |
| **Rotate structure** slider + angle box | Turns the whole drawing up to a full turn either way: −360° to 360°, with 0 in the middle. Positive is clockwise, negative counter-clockwise. Drag the slider, or type an exact angle in the box next to it (decimals are fine) and press Enter; the slider moves to exactly what you typed. The two always stay in sync. Letters always stay upright. |
| **Fit to view** | Re-centers and zooms so the whole drawing fills the canvas. |
| **Rotation 0°** | Snaps the whole-structure rotation back to zero. |
| **Reset all positions** | Undoes every manual move (drags and rotated selections) in the current view, returning to the automatic layout. |
| **Rotate selection**: degrees | Angle to turn the selected bases. Positive = clockwise, negative = counter-clockwise. |
| **Pivot** | *Attachment point*: turns the selection about where it joins the rest of the structure (for a stem, its closing base pair), so it swings like a hinge. *Center of the selection*: spins it in place. |
| **Rotate selected** | Applies the rotation. Each click adds another turn, and every rotation can be undone. |
| **Lock ring** / **Unlock ring** | Varna-style ring rotation. Ctrl/⌘+click an unpaired base of a loop to select its ring, then click **Lock ring** (a dashed circle marks it). Now dragging any helix attached to that ring swings it around the ring; dragging the closing stem swings the whole rest of the molecule. The ring keeps its size and its unpaired bases spread out evenly between the helices, which act as dividers. A helix stops when its neighbour's arc gets too tight, so nothing overlaps. The ring's own bases are locked. Each swing is one undo step. Click **Unlock ring** or press  Esc  when done. Single-sequence view only. |
| **Free tails** | How unpaired 5′/3′ ends are drawn: *Natural* (curves inward like a loop: on a single hairpin the 5′ and 3′ ends form an open loop at the base of the stem that mirrors the hairpin loop, never touching; with several stems each end curls toward the stems), *Flatten* (straight line), *L-shape* (one bond straight, then a 90° corner outward), *Perpendicular* (turns 90° outward). Changing the style clears any drags on the tail bases. In a complex it also shapes each strand's leftover ends. |

The **Manual positions** box on this tab holds **Reset all positions**; click its **(i)** for a short reminder of the drag gestures. They’re explained fully in section 5.

#### 4.3 Bases tab: letters and circles

Everything about the bases themselves is on one tab, in five boxes from top to bottom: **Letters**, **Circles**, **Color by base**, **Selected bases** and **Color from file**. Letter size and circle size are separate controls, so you can draw small letters in big circles or the other way round.

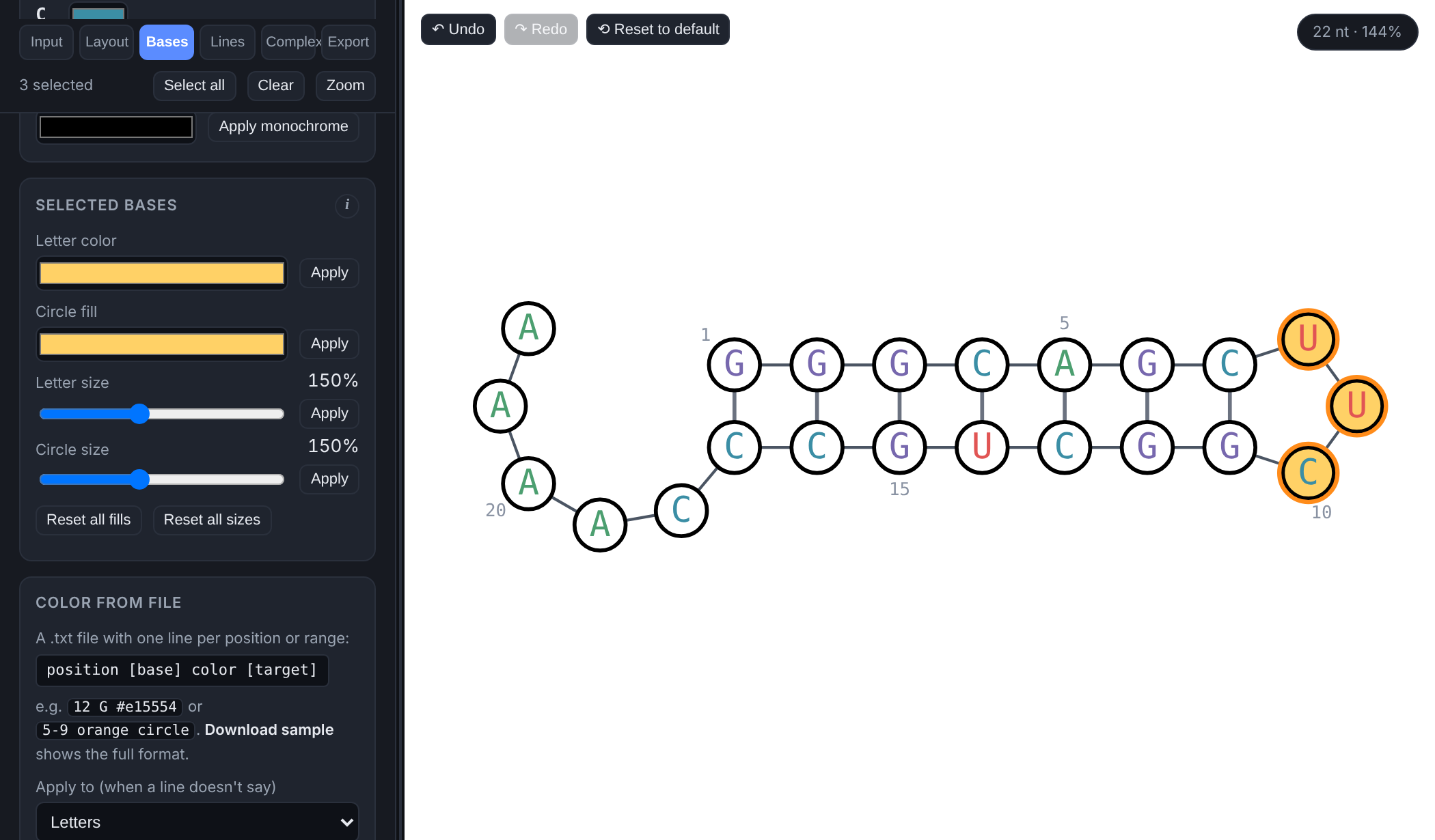

*Figure 3. Bases tab, scrolled to the Selected bases box. Circles on, numbering every 5 nt, and a custom circle fill applied to the three selected loop bases.*

| **Control** | **What it does** |
| --- | --- |
| **Letter size** slider | 8–60 px. Changes only the letters. Position numbers and 5′/3′ labels are sized relative to it. Circles keep their own size (below). |
| **Show base letters** | Turns the letters on or off (e.g. for circles-only figures). |
| **Bold letters** | Bold weight for all letters. |
| **Show base circles** | Draws a circle behind every base. |
| **Circle size (diameter)** slider | 6–120 px. Changes only the circles. Lines stop at the edge of whichever is bigger, the circle or the letter, so they never run into either. |
| **Circle frame** menu + swatch | Outline color: *Custom color* (use the swatch), *By base color* (identity palette), or *None*. |
| **Frame thickness** slider | 0.5–12 px outline width. |
| **Circle fill** menu + swatch | Inside color: *Custom color*, *None* (see-through), or *By base color*. |
| **Color by base** (A U T G C swatches) | The identity palette. Changes show up immediately on circles set to "By base color"; letters take the new palette when you click **Apply palette**. |
| **Apply palette** | Recolors **every** letter by base identity. |
| **Monochrome letter color** + **Apply monochrome** | Recolors every letter in one color. |

| **Control** | **What it does** |
| --- | --- |
| **Selected bases**: **Letter color** + **Apply** | Recolors only the selected letters. |
| **Circle fill** + **Apply** | Gives just the selected bases their own circle fill, overriding the global fill. |
| **Letter size** slider + **Apply** | Makes the selected letters bigger or smaller (50–300%), relative to Letter size. Their circles don’t change. |
| **Circle size** slider + **Apply** | Makes the selected bases’ circles bigger or smaller (50–300%), relative to Circle size. Their letters don’t change, and base circles are switched on if they were off. Very large sizes can touch neighbouring bases; drag them apart if needed. To size one base on its own, right-click it. |
| **Reset all fills** | Removes every per-base fill so all circles use the global fill again. |
| **Reset all sizes** | Returns every letter and circle to 100%. |
| **Color from file**: **Apply to** menu | Where colors from the file go when a line doesn't say: *Letters*, *Circle fill* or *Both*. |
| **Upload color file…** | Reads a .txt file of colors and applies them (see 4.3.1). A note under the buttons says how many bases were colored and lists any lines it skipped. |
| **Download sample** | Saves a ready-to-edit color file for whatever is open, with every position listed and colored by part (stems blue, loops orange, free ends gray), plus comments explaining each column. |

##### 4.3.1 Color file format

A plain .txt file, one base (or range) per line:  position [base] color [target] . Columns can be separated by spaces, tabs or commas. Lines starting with  #  are comments; a header line is ignored.

| **Column** | **What to put** |
| --- | --- |
| position | 1-based position ( 12 ) or a range ( 5-9 ). Inside a two-strand complex, prefix  A:  or  B:  to pick the strand ( B:12 ); no prefix means strand A. |
| base (optional) | The base letter at that position. Only used as a check: if it doesn't match the sequence, the app warns that the file may be for a different sequence. |
| color | Hex ( #e15554  or  #e15 ) or a color name ( red ,  orange ,  steelblue …). |
| target (optional) | letter ,  circle  or  both . Without it, the **Apply to** menu decides. Applying circle colors turns base circles on automatically. |

Example lines:  12 G #e15554  ·  5-9 orange circle  ·  B:3, #3b6fb6, both . Positions not listed keep their current color, so you can layer several files.

**Watch out Apply palette** and **Apply monochrome** overwrite any per-base letter colors you set earlier. Apply them first, then color your selections.

#### 4.4 Lines tab: backbone, rungs and numbering

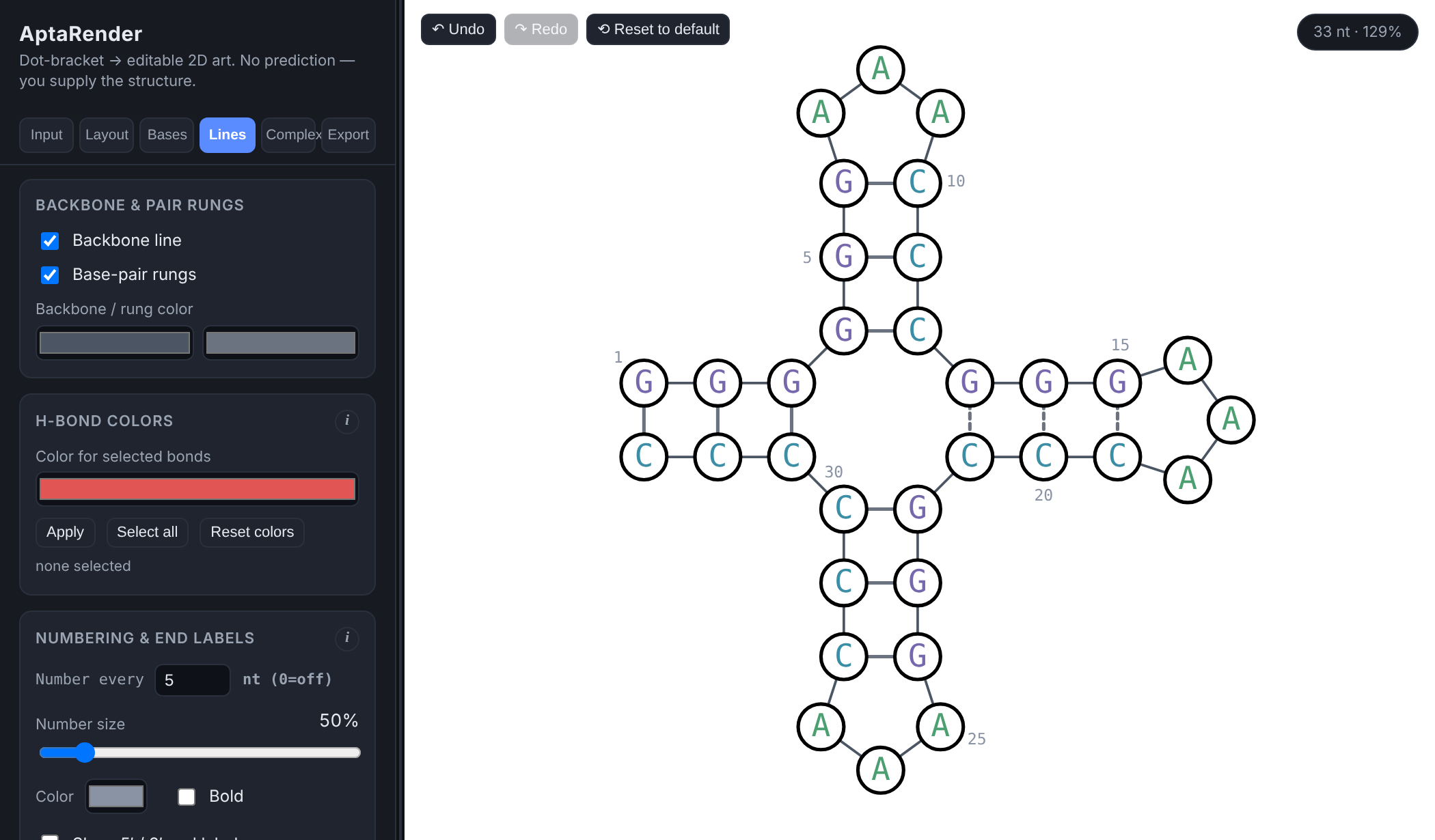

*Figure 4. Lines tab. The middle-right stem was written with [ ] so its rungs are dashed; numbering every 5 nt.*

| **Control** | **What it does** |
| --- | --- |
| **Backbone line** | Shows the line joining consecutive bases. |
| **Base-pair rungs** | Shows the lines between paired bases (solid for ( ), dashed for [ ] { } < >). |
| Two color swatches | Left: backbone color. Right: default rung color. |
| **Color for selected bonds** + **Apply** | Colors the selected rungs. Select a rung by **clicking it** on the canvas; **Shift**+click adds more. Selected rungs glow yellow. |
| **Select all** (bonds) | Selects every rung. |
| **Reset colors** | Returns every rung to the default rung color. |
| **Number every … nt** | Labels every Nth base with its position (0 = off). The first base is always labeled. Each number is placed beside its base in open space (outside loops, on the outer side of stems), never on top of a base or line; if it has to sit a little further out, a thin line connects it to its base. |

Individual rung coloring works on single sequences. In a complex, rungs keep the colors they were given on each strand's own sequence.

#### 4.5 Complex tab: two-strand hybridization

A **complex** draws two of your saved sequences together with a stretch of one pairing with a stretch of the other. You choose the region; nothing is predicted.

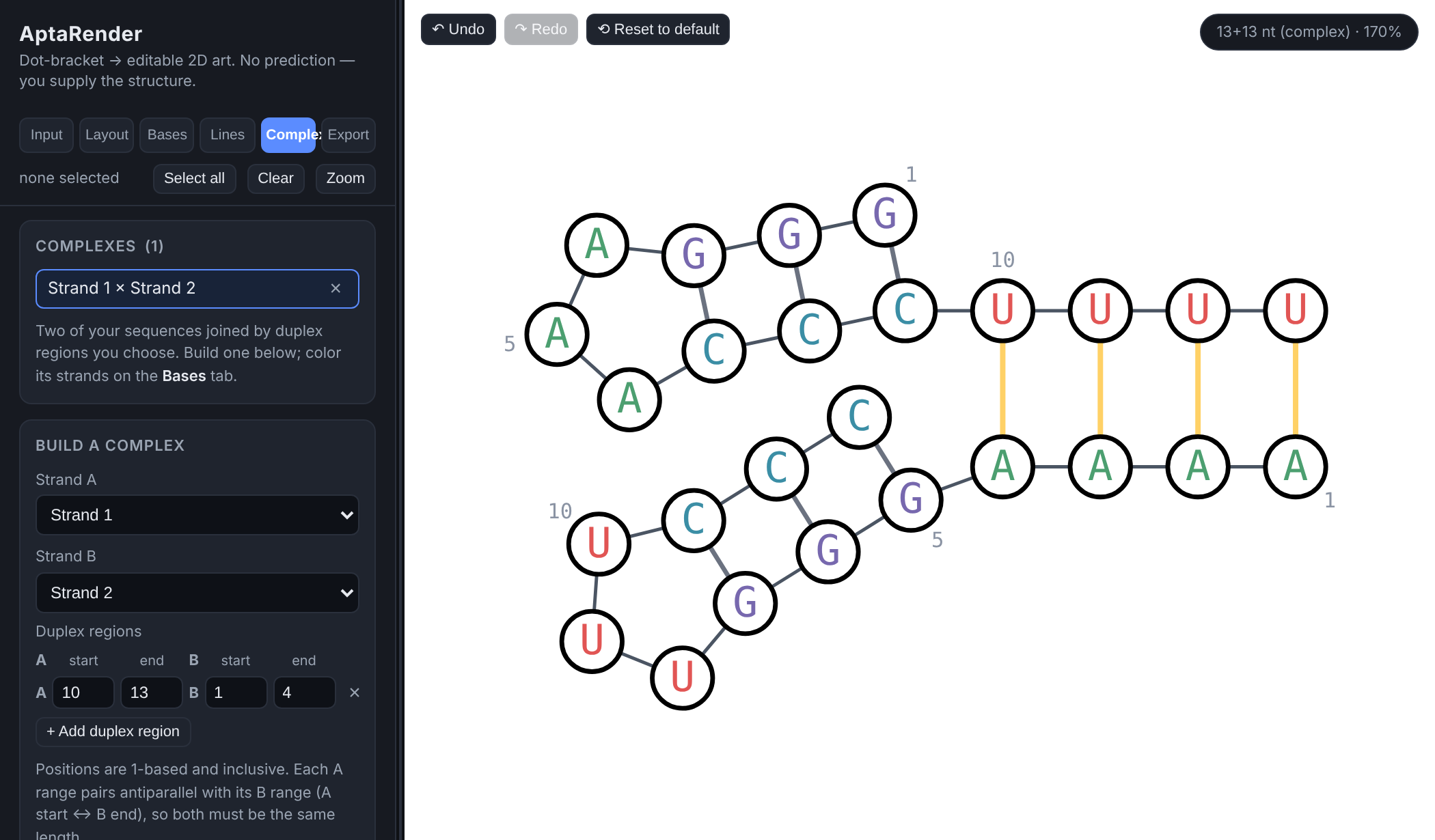

*Figure 5. A Replace-mode complex. The four yellow rungs are the inter-strand duplex.*

| **Control** | **What it does** |
| --- | --- |
| **Complexes** list | Your saved complexes. Click a name to open it, **×** to delete it. Side-by-side ones are tagged "· side". |
| **Strand A** / **Strand B** | Pick two different sequences from your list. Both need a valid sequence and structure first. |
| **A range** / **B range** | Start and end positions (1-based, inclusive) of the pairing region on each strand. Both ranges must be the same length. They pair antiparallel: A's start pairs with B's end. |
| **Mode: Replace** | Any existing pair in either strand that touches the region is broken, and the region is drawn as a fresh straight duplex. The rest of each strand hangs off its end. |
| **Mode: Side-by-side** | Strand A is left exactly as it is. Strand B is drawn alongside A's chosen arm, pushed out far enough to clear the rest of A. Needs at least 2 nt. If A's range has no pairs of its own you'll get a hint that you probably meant to swap A and B. |
| **Build complex** | Creates the complex, opens it, and reports how many existing pairs were broken. |
| **Duplex rung color** | Color of the inter-strand rungs. |

##### Working inside a complex

- Bases, Lines, Layout, Export, selection, dragging, rotation, numbering and undo all work in a complex exactly as they do on a single sequence.
- Colors you apply to bases in a complex are saved **on the two source sequences**, so they also appear when you open those sequences on their own, and vice versa.
- Dragged positions and the view (zoom, rotation) are saved **with the complex**.
- To leave the complex, click any sequence name on the Input tab.
- If you later edit a source sequence's structure, reopening the complex redraws it from the new structure.

#### 4.6 Export tab

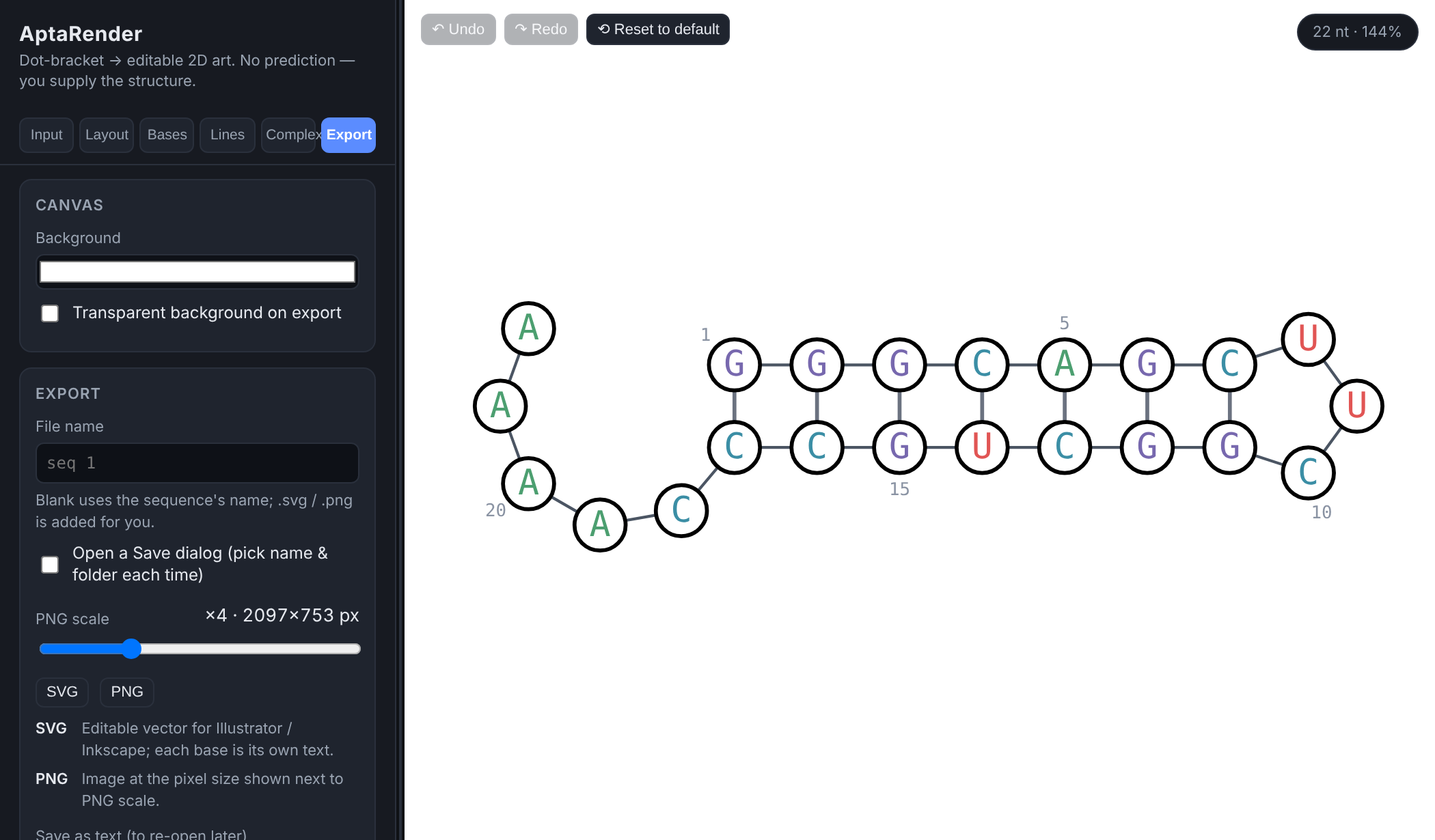

*Figure 6. Export tab, showing the PNG pixel size for the current scale.*

| **Control** | **What it does** |
| --- | --- |
| **Background** | Canvas color, also used as the export background. |
| **Transparent background on export** | Leaves the exported SVG or PNG without a background, for placing on colored slides or posters. |
| **PNG scale** slider | 1×–12×. The label shows the resulting PNG size in pixels. Use 4× or higher for print. |
| **File name** | Name for the downloaded file. Leave it blank to use the sequence's (or complex's) name; .svg or .png is added for you, and characters file systems don't allow are replaced. |
| **Open a Save dialog** | In Chrome and Edge, opens the system Save dialog for each export so you can rename the file and choose the folder. Not available in Safari or Firefox; in Safari, turn on Settings → General → File download location → Ask for each download to get the same effect. |
| **SVG** | Downloads an .svg file: fully editable vector art (every letter is its own text element) for Illustrator, Inkscape or Affinity. |
| **PNG** | Downloads a .png file at the chosen scale. |

Exports contain exactly what's on the canvas (colors, rotation, dragged positions, numbering) cropped to the drawing with a small margin. Selection rings and highlight glows are left out, and zoom and pan don't matter.

### 5. Working on the canvas (mouse)

#### 5.1 Selecting

| **Do this** | **Result** |
| --- | --- |
| **Click** a base | Selects only that base. Click it again to deselect. |
| **Shift** + click a base | Adds it to, or removes it from, the selection. |
| **Ctrl/⌘** + click a base | Adds or removes its whole **part** (see 5.3). |
| **Shift** + drag on empty canvas | Draws a dashed box. Every base inside is added to the selection. |
| **Click** a rung | Selects that base pair for bond coloring (Lines tab). **Shift**+click adds more. Single sequences only. |
| **Click** empty canvas | Clears the base and bond selection. |

#### 5.2 Moving and viewing

| **Do this** | **Result** |
| --- | --- |
| **Drag** a base | Moves its whole **branch**: for a paired base, the stem and everything it encloses; for an unpaired base, just that base. The rest stays put. |
| **Alt/Option** + drag a base | Moves only that single base. |
| **Ctrl/⌘** + drag a base | Moves only its **part**: just the helix, just the loop ring, or just the free end. |
| **Drag a selected base** (when 2 or more are selected) | Moves the whole selection together, even across both strands of a complex. |
| **Drag** empty canvas | Pans the view. |
| **Scroll wheel / trackpad scroll** | Zooms in or out around the pointer (5% to 3000%). |

**Tip** Every move is recorded as one undo step, and **Reset all positions** (Layout tab) clears them all at once.

#### 5.3 What is a "part"?

Parts let you grab a natural piece of the structure without selecting bases one by one. AptaRender works them out from the structure automatically:

- **Helix**: click a paired base and you get the unbroken stack of base pairs it belongs to (both strands of the stem), but not the loop at its tip.
- **Loop ring**: click an unpaired base inside a loop and you get all the unpaired bases of that loop (hairpin, internal or multiloop), skipping any stems branching off it.
- **Free end**: click an unpaired base outside every pair and you get its whole unpaired run, such as the 5′ or 3′ tail or a linker between stems.

In the Hairpin example, Ctrl/⌘+clicking any of the three loop bases selects exactly those three (that's how Figure 3's highlight was made), and Ctrl/⌘+dragging an A in the tail moves only the tail.

### 6. Keyboard shortcuts cheat sheet

On a Mac, use **⌘ Cmd** wherever Ctrl is listed and **Option** for Alt.

| **Shortcut** | **Action** |
| --- | --- |
| Ctrl + Z  /  ⌘ + Z | Undo |
| Ctrl + Y ,  Ctrl + Shift + Z  /  ⌘ + Shift + Z | Redo |
| Shift  + click base | Add/remove base from selection |
| Shift  + drag background | Box select |
| Ctrl/⌘  + click base | Add/remove its part from selection |
| Ctrl/⌘  + drag base | Move only its part |
| Alt/Option  + drag base | Move only that base |
| Esc | Unlock a locked ring (ring rotation) |
| Shift  + click rung | Add/remove bond from bond selection |
| Scroll | Zoom around pointer |
| Double-click sequence name | Rename sequence |

Undo and redo shortcuts pause while you're typing in a text or number box, so the box's own undo works there. Click on the canvas first to use AptaRender's undo.

### 7. Recipes for common figures

#### 7.1 A clean publication figure

1. Enter the sequence and structure on **Input** and click **Render**.
2. **Bases**: set **Letter size**, then **Apply palette** (color by base) or **Apply monochrome**.
3. **Bases** (Circles box): tick **Show base circles**, set **Circle size**, and choose frame and fill (e.g. black frame, white fill, 2 px).
4. **Lines**: set **Number every** to 10.
5. **Layout**: rotate the whole structure so it reads well; drag any crowded branch into a clearer spot.
6. **Export**: tick **Transparent background**, set **PNG scale** to 4× or more, and download **SVG** and/or **PNG**.

#### 7.2 Highlight a loop or binding site

1. Ctrl/⌘+click one base of the loop to select the whole ring, or Shift+drag a box around the bases you want.
2. **Bases** tab → **Selected bases** → pick a color under **Circle fill** → **Apply**.
3. Optionally, in the same box, pick a **Letter color** → **Apply**, e.g. white letters on a dark fill.
4. Click empty canvas to clear the selection and check the result.

#### 7.3 Swing a crowded stem out of the way

1. Click a base pair at the bottom of the stem, then Shift+drag a box over the rest of that stem and its loop. Or simply drag a base of the stem: the whole branch moves with it.
2. To turn it instead of moving it: **Layout** tab → **Rotate selection**, enter an angle (try 30 or −30), set **Pivot** to *Attachment point*, click **Rotate selected**. Repeat until it looks right.
3. Press **Zoom** in the selection bar to inspect it up close, then **Fit to view** to see the whole figure again.

#### 7.3b Re-angle a helix around a loop (Varna-style)

1. Ctrl/⌘+click any unpaired base in the loop the helix is attached to. Its whole ring is selected.
2. **Layout** tab → **Lock ring**. A dashed circle marks the locked ring.
3. Drag a base of the helix in a circle around the ring. It slides along the ring while the ring's unpaired bases re-spread on either side.
4. To swing everything else instead, drag the stem that closes the ring.
5. Click **Unlock ring** (or press  Esc ) and **Fit to view**.

#### 7.3c Color by your own data (DMS, mutations…)

1. Open the sequence, then **Bases** tab → **Color from file** → **Download sample**.
2. Edit the color column (or delete lines you don't need, and use ranges like  20-25 ). Keep the base column as a check that the file matches the sequence.
3. Choose **Apply to** (letters, circle fill, or both), then **Upload color file…**.
4. Read the note under the buttons for anything skipped or mismatched.

#### 7.4 Show two strands hybridizing

1. On **Input**, add both strands as separate sequences (use **+ Add**, or import a FASTA with both) and **Render** each one so it has a valid structure.
2. Go to **Complex**, choose **Strand A** and **Strand B**.
3. Enter the pairing ranges. Both must be the same length; remember A's first base pairs with B's last.
4. Choose **Replace** to draw the region as a clean duplex, or **Side-by-side** to keep A's own fold untouched and lay B along it.
5. Click **Build complex**, then style and export it like any other figure. Set the **Duplex rung color** to make the inter-strand pairs stand out.

### 8. Saving, undo, and where your data lives

- **Saving is automatic.** Everything (sequences, structures, colors, positions, complexes, styles, panel width) is stored in your browser as you work and comes back when you reopen the page.
- **It stays on this computer, in this browser.** Nothing is uploaded and nobody else can see it. A different browser or computer won't have it, and clearing your browser's site data deletes it. Export an SVG of anything you want to keep.
- **Undo goes back up to 100 steps** and covers rendering, colors, every style control, drags, rotations, adding/deleting/renaming sequences, and building/deleting complexes.
- Switching which sequence is open, changing the selection, and panning/zooming aren't undo steps, so Ctrl+Z won't jump you around.

### 9. Limitations and troubleshooting

| **Situation** | **What to know / do** |
| --- | --- |
| Circles didn’t change size with the letters | Since version 2.3, letter size and circle size are separate. Use **Circle size** in the Bases tab (or under **Selected bases** for just some bases). |
| My custom colors disappeared | **Render**, **Reset to default**, **Apply palette** and **Apply monochrome** reset colors. Press Ctrl/⌘+Z. |
| A branch looks misplaced after changing Free tails | Tail style re-lays out the drawing. Drags on the tail bases are cleared, but moves on stems and loops are kept. Use **Reset all positions**, then re-adjust. |
| Pseudoknots | Crossing pairs aren't supported. All brackets must nest. Use [ ] { } < > only to mark non-WC pairs. |
| Structure prediction | Not provided by design. Supply the structure from your own folding tool or data. |
| Bond coloring in a complex | Rung click-selection works on single sequences only. Color bonds on each strand's own sequence first; they carry into the complex. |
| More than two strands / several duplex regions | A complex is two strands with one contiguous pairing region in this version. |
| Hit the 20-sequence limit | Delete a sequence you no longer need (its complexes go with it). |
| Something else | Use the contact link at the bottom of the Export tab. |
